# All ribosomal RNAs and 45S spacers from humans to worms are packed with organism-specific motifs whose other copies are found predominantly in numerous nervous system genes including many associated with human disorders

**DOI:** 10.1101/2024.12.10.627812

**Authors:** Isidore Rigoutsos, Stepan Nersisyan, Eric Londin, Iliza Nazeraj, Bonnie Dong, Anastasios Vourekas, Phillipe Loher

## Abstract

The nucleotide sequences of ribosomal RNAs (rRNAs) and the spacers of 45S are tuned to fulfill optimally their respective roles during ribosome biosynthesis and function. We report that these sequences satisfy additional genome-wide constraints in humans, mice, fruit flies, and worms. In all four organisms, the rRNAs and 45S spacers are densely packed with organism-specific nucleotide motifs with many additional identical copies throughout the genome. The human rRNAs and 45S spacers contain 1,723 motifs whose sequences are unique to the rRNAs/spacers of primates. These motifs have numerous additional exact intronic and exonic copies whose genomic placement is also unique to primates. Specific combinations of the motifs appear exclusively in 3,430 human nervous system and developmental genes, including 1,046 risk genes for autism, schizophrenia, and bipolar disorder. RNA/RNA crosslinking experiments show that the rRNA/spacer motifs are contact points for rRNA-mRNA and mRNA-mRNA heteroduplexes. RNA binding protein (RBP) assays show that these motifs are also in the binding sites of 113 RBPs. RNA sequencing reveals that rRNAs and spacers produce endogenous small non-coding RNAs (sncRNAs) that carry the same primate-specific motifs. Lastly, the motifs’ intergenic and intronic copies overlap 131 GWAS polymorphisms associated with neuropathologies (p-val<3.9e-12). The findings suggest that the motifs facilitate RNA/RNA and RBP/RNA interactions that are affected by polymorphisms and modulated by rRNA- and spacer-derived sncRNAs carrying the same motifs. Our study also genetically links for the first time rRNAs and 45S spacers to autism and other typically human neurological disorders. Mutation panels based on these motifs can lead to new molecular diagnostics for these disorders, whereas snRNAs carrying these motifs can serve as drugs or potential therapeutic targets.

**Graphical abstract:** 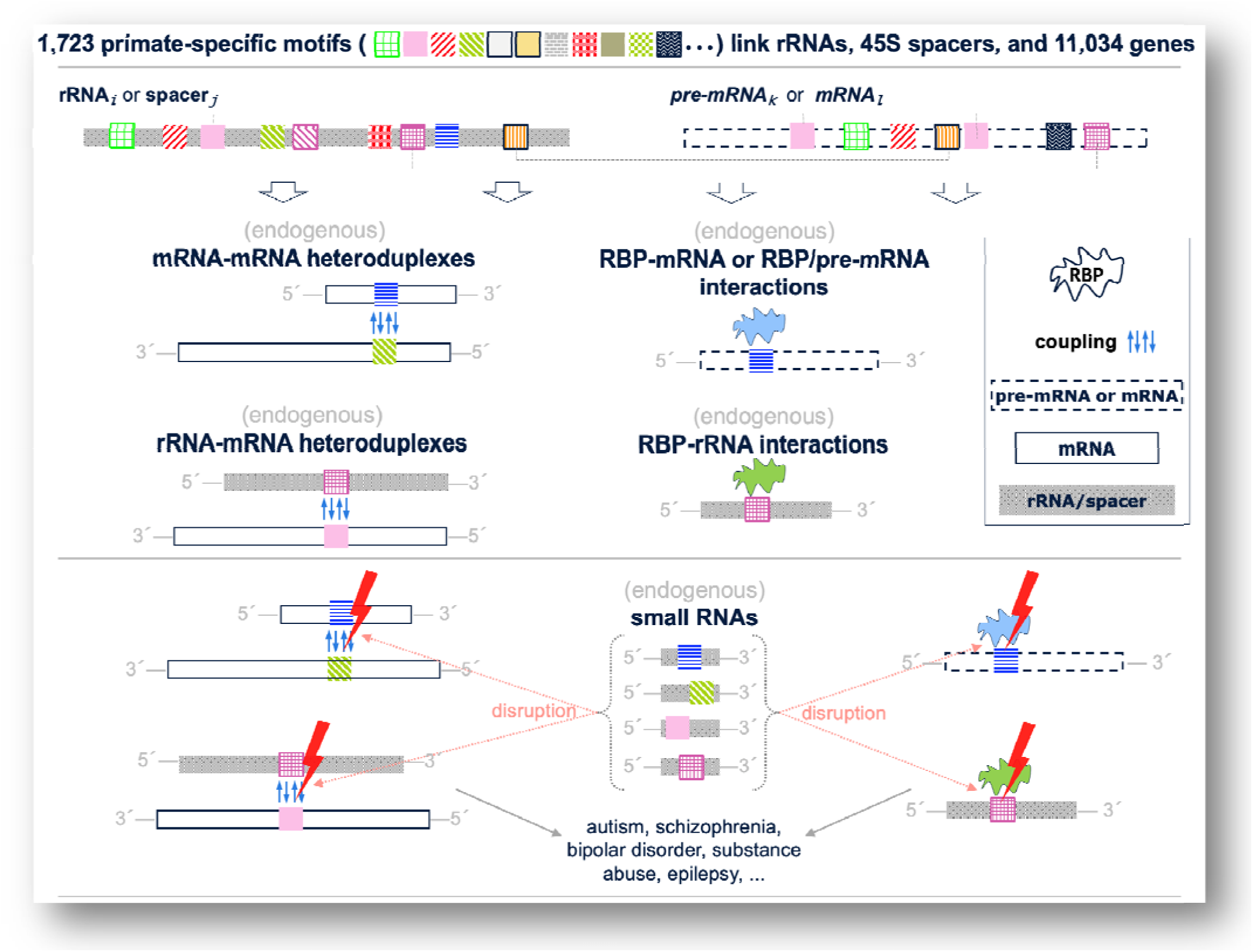

The human ribosomal RNAs and the 45S spacers share primate-specific motifs with thousands of nervous system genes, including risk genes for autism and other neurological disorders. Similar links between rRNAs/spacers and nervous system genes exist in mice, fruit flies, and worms but are achieved through motifs that are unique to each organism. In humans, the motifs are at the contact points of endogenous gene-gene and rRNA-gene heteroduplexes, as well as the binding sites of RNA binding proteins (RBPs) in genes and rRNAs/spacers. Endogenous small RNAs carrying these shared motifs modulate these endogenous interactions and are disrupted by mutations at the motifs’ genomic copies. Mutation panels based on these motifs can lead to new diagnostics for autism and other disorders, while the small RNAs carrying the motifs can serve as drugs or potential therapeutic targets.

## INTRODUCTION

We previously reported findings on a class of nucleotide (nt) motifs with many identical copies that are conserved across the genome, at least one of which is in an mRNA^1^. Requiring that a motif be present in an mRNA ensures that it is part of at least one transcript linked to protein production. To ensure statistical significance, we only considered motifs comprising at least 16 consecutive nts with 30 or more intergenic or intronic copies. We dubbed these motifs “pyknons,” from the Greek word for “dense, serried.”

Those studies were carried out on early assemblies of the human and mouse genomes that were incomplete and less well annotated compared to what is now available to us. Nonetheless, we were able to show that pyknons are genome-specific^1^, are placed non-randomly in mRNAs^1^, appear in the mRNAs of genes belonging to specific biological processes^1^, co-localize with regions that produce piRNAs^2^, and capture *functional* conservation between human and mouse without *sequence* conservation^2,3^. A subsequent independent study of the *A. thaliana* genome led to similar findings^4^.

In follow-up work, others and we found that RNAs containing these motifs have characteristic tissue-specific and disease-specific expression profiles in various settings^5^, including colon cancer^6,7^. For the long noncoding RNA (lncRNA) N-BLR, we showed that it facilitates the epithelial to mesenchymal transition in colon cancer cells by using a pyknon near its 3’ terminus to decoy a microRNA (miRNA) target^5^. Another group showed that the DNA methyltransferase DNMT1A, in addition to binding DNA, also binds RNAs at sites enriched for pyknons^8^. More recently, we reported that the mRNAs whose levels *decrease* from the zygote to the blastocyst contain vastly different pyknons than those whose levels *increase* during the same period^9^.

In the current study, we build on these earlier efforts^1–3,5,7,9^ by leveraging the availability of the telomere-to-telomere human genome assembly^10^, which added more than 200 million nucleotides to the human genome including all copies of the 45S cassette, and the sequences around centromeres and telomeres. Using the first instance of a human genome assembly that is essentially complete, we sought to answer the following question: “Do the specifics of a genome’s architecture bring about regulatory linkages between the genome’s non-coding and protein-coding regions, including regions with known disease associations?” Using a statistically rigorous approach, we examined the human genome “bottom-up,” from its nucleotide sequence towards its annotated regions, including rRNAs, 45S spacers, regions coding for protein-coding genes or small non-coding RNAs (sncRNAs), and risk variants. We analyzed the sequences of the two mitochondrial rRNAs, the four nuclear rRNAs (5S, 18S, 5.8S, and 28S), and the four transcribed spacers of 45S and studied their linkages to genes and various conditions and disorders. Lastly, we extended the analysis to three more organisms: the mouse, the fruit fly, and the worm.

## METHODS

### Generating the pyknons

We computed the pyknons in the T2T human genome assembly^11^ by generating 16-mers, intersecting them with the 96,441 mRNAs (22,511 protein-coding genes) of Rel. 105 of ENSEMBL, and removing redundancies as previously described^1^. See Supplemental Methods for more information.

### Reference rRNAs

We obtained the reference sequences of 5S, 12S, 16S, and 45S from GenBank. We obtained the sequences of genes (mRNAs and pre-mRNAs) from ENSEMBL. See Supplemental Methods.

### Repeats

We downloaded RepeatMasker annotations for the T2T genome from the UCSC Table Browser (table ID: “hub_3671779_t2tRepeatMasker”). We used an exhaustive, brute-force search to find all appearances of the 1,723 pyknons within the known repetitive sequences.

### Enrichment analyses

We determined enrichments for GO terms, KEGG pathways, and The Alliance of Genome Resources diseases with ShinyGO^12^. See Supplemental Methods.

### Searches for syntenic sequences

For each pyknon instance within a gene’s 5’-UTR, CDS, 3’-UTR, or intron, we determined its syntenic counterpart in other organisms as follows: we examined the sequence immediately upstream and downstream of the pyknon’s copy and extended it to include any additional adjacent instances; we flanked the potentially extended sequence with 100 nts on each side; finally, we sought the “flank+pyknon copy/-ies+flank” in the target genome at the location of the orthologous gene ±100 kb. See Supplemental Methods.

## RESULTS

We first identified 9,961,431 unique 16-mers with ≥ 30 identical copies across the T2T genome. We confirmed that these 16-mers are statistically significant through a Monte Carlo simulation that estimated the probability of a 16-mer having ≥ 30 identical copies by chance in the T2T genome to be ≤ 1.7*10^-6^. We intersected the initial collection of 16-mers with the 96,441 ENSEMBL mRNAs, removed redundancies using a previously described approach (see Supplemental Methods), and identified the pyknons of T2T, which we then intersected with the sequences of the six human rRNAs and the four transcribed spacers of 45S.

### The four nuclear rRNAs and the four spacers of 45S are densely packed with pyknons

We found 1,723 pyknons in the encodings of the six rRNAs and four 45S spacers: 843 pyknons appear only in sense, 848 in antisense, and 32 in both orientations (**Supp. Table S1A**). Most of the pyknons are found in the four nuclear rRNAs and the transcribed spacers of 45S, which are densely packed with pyknons in both sense and antisense (**Supp. Figures S1-S2**). **Figure 1** shows secondary structures for two rRNAs (18S, 28S) and two spacers (ITS1, ETS2) – the structures of the two RNAs were determined by R2DT^13^ and those of the two spacers by RNAfold^14^. In each structure, we colored the regions covered by pyknons. We will discuss the different color codes in the next section. Most of the shown pyknon-covered regions span more than 16 nts and result from distinct pyknons that overlap partially but not entirely (see Supplemental Methods). **Supp. Table S1B** lists the pyknons in each rRNA and transcribed spacer and their respective location and orientation. **Supp. Table S1C** lists the pyknons’ genomic counts, which range between 30 and 174,822.

**Figure 1.**
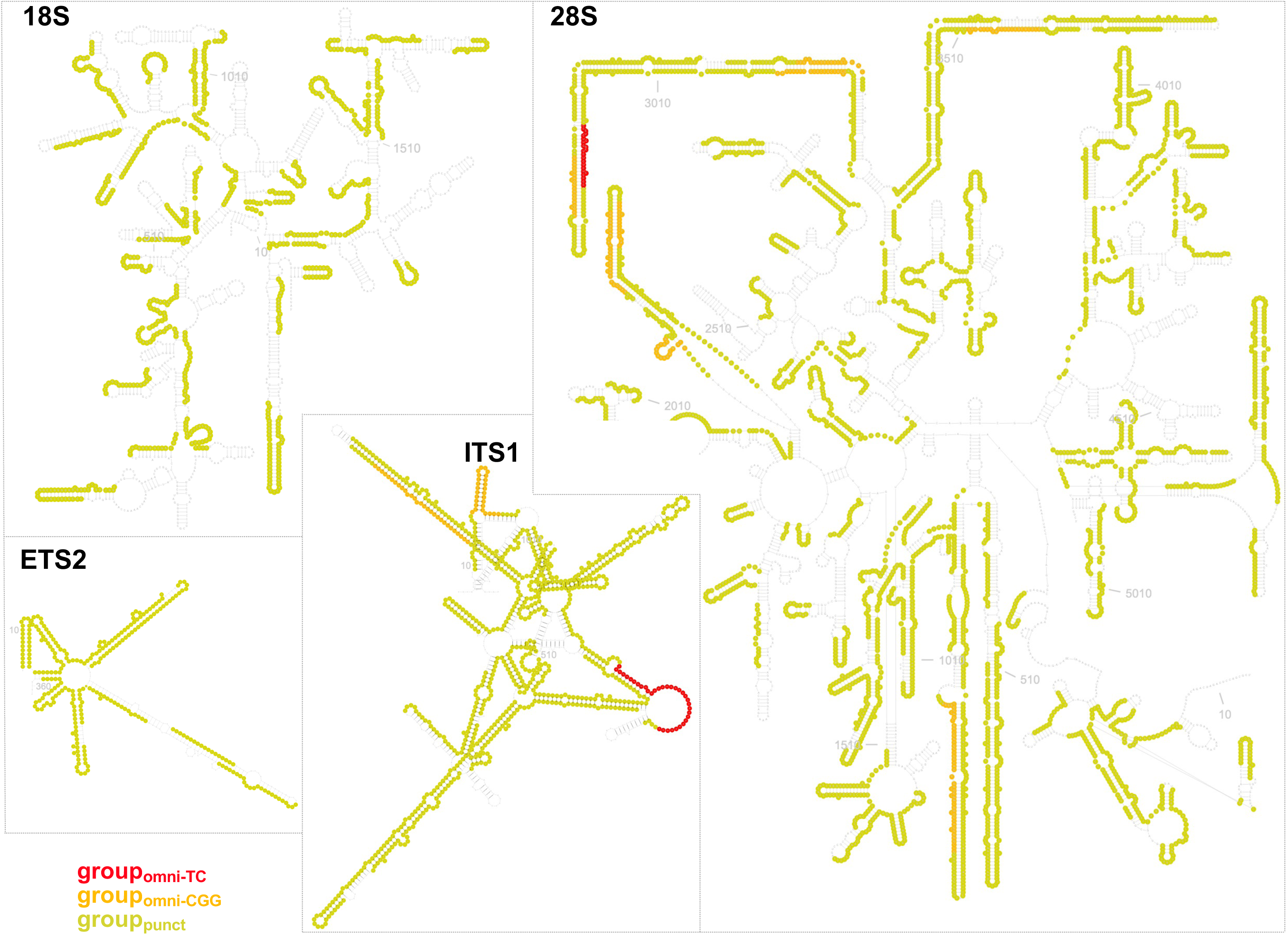
Secondary structures of 18S and 28S rRNAs and the ITS1 and ETS2 spacers. The pyknon-covered regions are highlighted. Red: instances of sequences from group_omni-TC_. Orange: instances of sequences from group_omni-CGG_. Yellow: instances of sequences from group_punct_. The styles of secondary structures were modified in RNAcanvas^69^.

Given the recurring nature of pyknons in the genome, we examined whether any of the 1,723 pyknons in rRNAs and spacers can be found within known repeat elements annotated by RepeatMasker (see Methods). Almost all pyknons – 1,663/1,723 or 96.5% – are present in long interspersed nuclear elements (LINEs) in sense or antisense orientation (**Supp. Table S1D**). The LINE-1 elements harbor over 91% (1,578/1,723) of the pyknons, followed by LINE-2 elements (1,340/1,723 or 78%). Importantly, most rRNA/spacer pyknons appear in annotated LINEs as *isolated islands*, 16 nts in width or slightly longer: in other words, the rRNA/spacer pyknons do not form long continuous blocks within these repeats. Thus, the presence of pyknons in rRNAs and spacers is non-trivial and does not follow from their presence in LINEs.

### The pyknons found in the rRNAs and the transcribed spacers of 45S form three groups

Looking at **Supp. Table S1C,** we can divide the rRNA/spacer pyknons into three groups based on their genomic counts and pairwise sequence similarity. The first group comprises the pyknon TCTCTCTCTCTCTCTC or “(TC)_8_,” its reverse complement, and their sequence variants. The second group includes the pyknon GCGGCGGCGGCGGCGG or “G(CGG)_5_,” its reverse complement, and their variants. The third group consists of all remaining pyknons. We identified the sequences of the (TC)_8_ and G(CGG)_5_ groups using an iterative procedure (see Supplemental Methods) that allowed members of the (TC)_8_ and G(CGG)_5_ groups to “attract” other pyknons among the 1,723 as long as their respective sequences were within a Levenshtein distance of 1. By design, the pyknons in the three groups are present in sense or antisense in rRNAs or spacers. To simplify the subsequent analyses, we added to each group the reverse complement of its pyknons, removed duplicates, and created the final three collections with 34, 141, and 2,635 sequences, respectively (**Supp. Table S1E**).

In what follows, we will refer to the 34 sequences as “group_omni-TC_” and the 141 sequences as “group_omni-CGG_”. The modifier “omni” acknowledges the genomic omnipresence (high genomic counts) of the respective sequences, as evidenced by **Supp. Tables S1C-D**. We will refer to the 2,635 sequences as “group_punct_” with the modifier “punct” highlighting the respective sequences’ considerably lower copy numbers and their punctate presence in protein-coding genes, as evidenced by **Supp. Table S1E**. Note that the pyknon (TC)_8_, which leads group_omni-TC_, has the highest number of genomic copies (174,822). On the other hand, the pyknon G(CGG)_5_, which leads group_omni-CGG_, appears in the mRNAs of the highest number of genes (540). In **Figure 1**, we used “red,” “orange,” and “yellow” to mark the instances of the sequences belonging to group_omni-TC_, group_omni-CGG_, and group_punct_, respectively.

### The (TC)_8_ and G(CGG)_5_ pyknons are very recent evolutionary insertions to 28S rRNA

(TC)_8_ is sense to 28S rRNA at one location (starting at position 3052) and antisense to ITS1 at three locations (starting at positions 191, 193, and 195, respectively). G(CGG)_5_ is sense to 28S at nine locations (starting at positions 849, 852, 870, 873, 876, 2153, 2156, 2159, and 3489, respectively) and antisense to ITS1 at one location (starting at position 741). The ten instances of (TC)_8_ and G(CGG)_5_ are located in the so-called “expansion segments” of the 28S rRNA^15,16^. When one compares humans to distant organisms, e.g., rodents^15^, the expansion segments are easy to locate thanks to their large size. Adding to these comparisons organisms that are phylogenetically closer to humans, e.g., non-human primates, permits the following additional observations.

**Figure 2** shows a multiple sequence alignment of 28S rRNA sequences from 20 organisms, spanning the spectrum from *H. sapiens* to *C. elegans*. The expansion segments appear as variable-size gaps in these sequences. However, in the sequences of the six primates, the gaps are much smaller, especially when comparing *H. sapiens* to chimp (*P. troglodytes*) and gorilla (*G. Gorilla*), five and seven million years away, respectively. The differences between humans, chimps, and gorillas, on the one hand, and all other primates and non-primates, on the other, correspond to the *insertions* of the ten instances of (TC)_8_ and G(CGG)_5_. At 15-20 million years away, in the 28S rRNA sequence of *N. leucogenys* (white-cheeked gibbon), only a single instance of G(CGG)_5_ remains. A notable exception is the presence of a single G(CGG)_5_ copy in the 28S rRNA from the odontocete *T. truncatus*, the common bottlenose dolphin.

### The pyknons found in the rRNAs/spacers have exact copies in sense or antisense in the pre-mRNAs of one-half of all protein-coding genes

We sought sense or antisense copies of the sequences in group_omni-TC_, group_omni-CGG_, and group_punct_ in the genomic spans of all 22,511 protein-coding genes. We found that the unspliced transcripts of 11,034 (49.0%) genes contain in sense or antisense one or more of the pyknons we find in rRNAs/spacers (**Supp. Table S2A**). This is a notable event because each of these pyknons is statistically significant.

### Most intronic and exonic instances of the pyknons from rRNAs/spacers are recent evolutionary insertions into the genomes of primates

Given that the 28S rRNA copies of (TC)_8_ and G(CGG)_5_ are not conserved beyond primates (**Figure 2**), we hypothesized that a similar absence of conservation must extend to all the pyknons we find in rRNAs/spacers and all their copies in rRNAs, spacers, and protein-coding genes.

• The sequences of the human 45S cassette pyknons are not conserved in the 45S cassettes of other organisms: We focused on the 1,692 pyknons found in the 45S cassette (2,754 unique sense or antisense sequences) and sought them in the 45S cassettes of five non-human primates, mouse, fruit fly, and the worm. We found that the number of human pyknons that are identically present in the 45S cassette of these six organisms decreases with phylogenetic distance. Specifically, of the 2,754 sequences we find in the human 45S cassette only a subset is present in *P. troglodytes* (1,571), *G. gorilla* (1,423), *P. abelii* (1,141), *M. mulatta* (962), *N. leucogenys* (930), *M. musculus* (418), *D. melanogaster* (58), and *C. elegans* (48), respectively. The number of sequences that remain in each organism’s cassette is inversely proportional to the organism’s phylogenetic distance from humans.
• The genic copies of human rRNA/spacer pyknons are not conserved in other organisms: We also analyzed all 31,319 exonic and intronic instances of the 1,723 rRNA/spacer pyknons, in sense or antisense, across the 11,034 protein-coding genes that contain them (**Supp. Table S2A**). We flanked each pyknon instance with 100 nts on each side, identified its syntenic counterparts in the other 19 genomes, and created the corresponding multiple sequence alignment. We computed a conservation score for the pyknon span (see Supplemental Methods) and separately for each flanking region. **Figure 3A** shows representative multiple-sequence alignments of syntenic regions from four genes, GALNTL6, AR, LMBR1, and GADL1, across several organisms. Note how (TC)_8_ is present in the human ortholog but only in some primates. **Supp. Figure S3A** shows several more alignments. **Figure 3B** highlights the difference between the conservation scores of the pyknon span and flanking regions, respectively. Note how ∼75% of all pyknon instances in 3’-UTRs and introns are less conserved than the nucleotide segments immediately flanking the pyknon. In 5’-UTRs and CDSs, the conservation percentages are lower but still above 50% and highly statistically significant. **Figure 3C** shows the average number of nucleotides (i.e., non-gaps) corresponding to the pyknon portion of the multiple sequence alignments for the intronic copies and the different genomes. All non-primates show a striking absence of intronic copies for the 1,723 pyknons or their antisense, indicating that most of these copies are insertions in the genomes of primates. **Supp. Figure S3B** shows counterpart plots for 5’-UTRs, CDSs, and 3’-UTRs.

**Figure 2.**
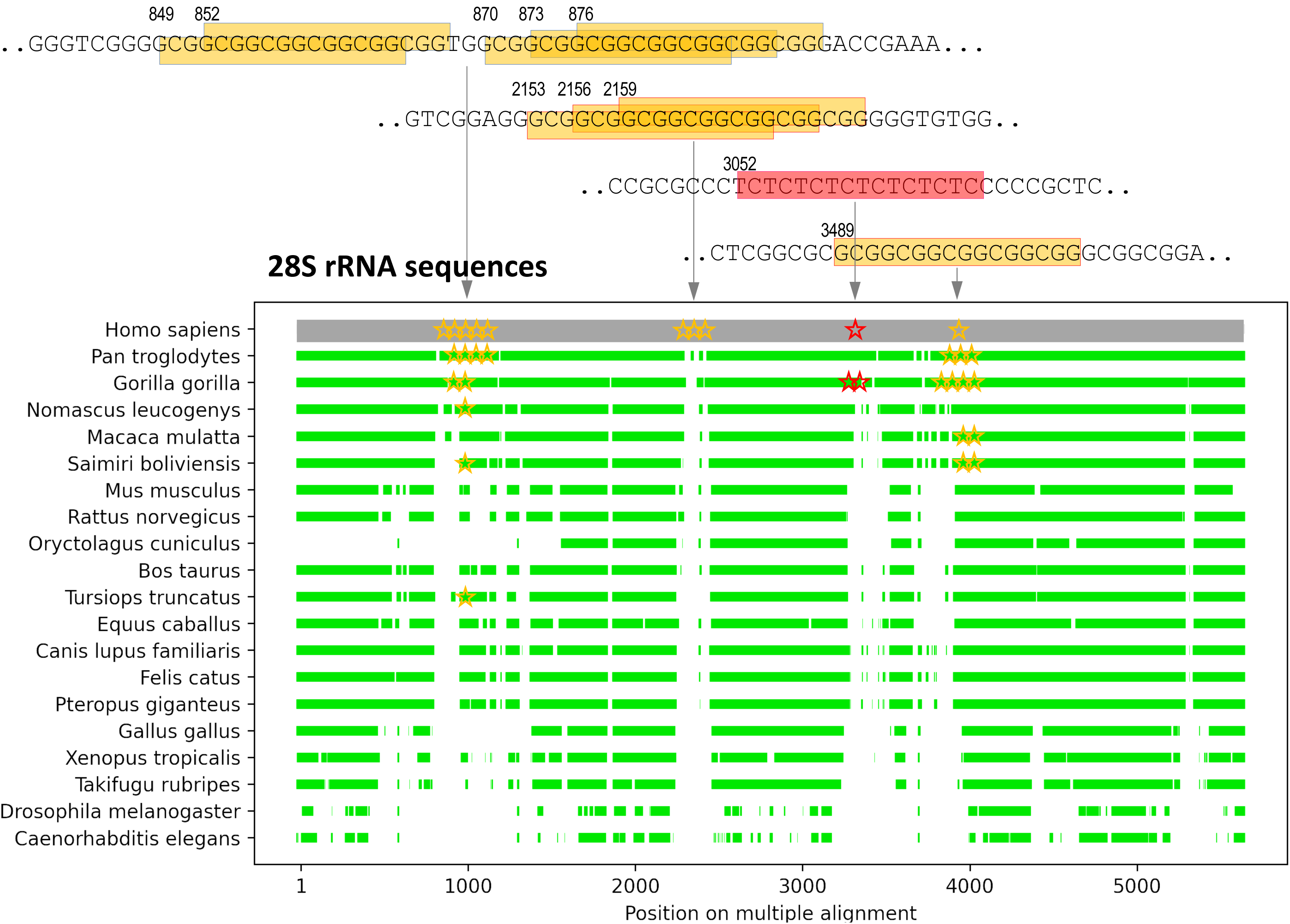
Multiple sequence alignments of 28S rRNA sequences from twenty organisms show prominently the existence and location of the previously-reported expansion segments of 28S rRNAs. A position of the alignment is colored in green if the corresponding 21-nt symmetric window has ≥ 75% of its nts matching the human sequence. Red and orange stars show the locations of the (TC)_8_ and G(CGG)_5_ pyknons, respectively. Note how in primates, where the expansion segments are established, the differences revolve around the insertions of the (TC)_8_ and G(CGG)_5_ pyknons.

**Figure 3.**
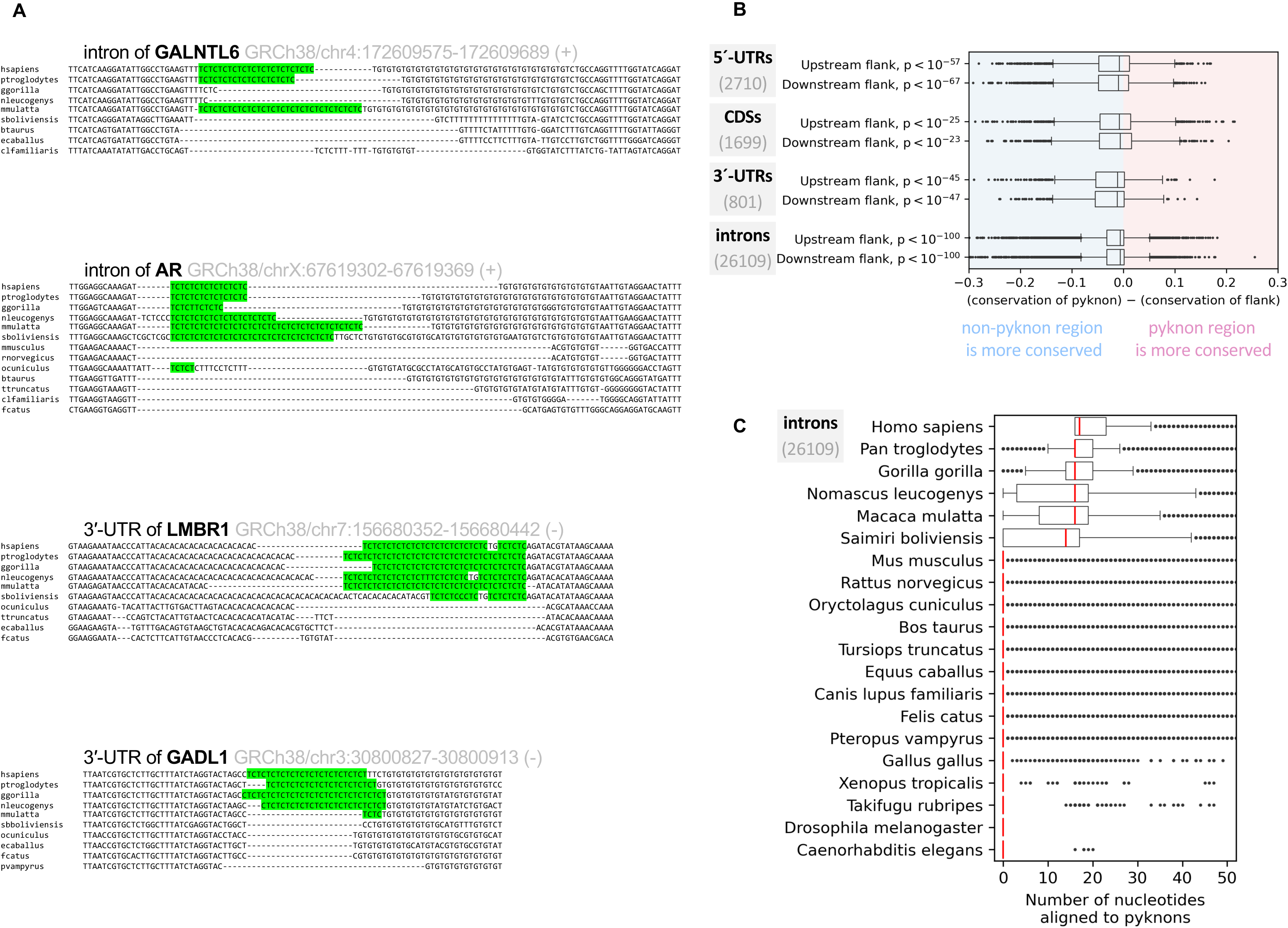
Conservation of group_omni-TC_, group_omni-CGG_, and group_punct_ instances across organisms. A: Multiple sequence alignments of syntenic regions from four representative genes. The human sequence contains an instance of (TC)_8_ from group_omni-TC_. The examples show a lack of conservation in organisms that are phylogenetically distant from humans. B: Boxplots showing the level of conservation of all exonic and intronic instances of the 2,810 sequences from group_omni-TC_, group_omni-CGG_, and group_punct_ instances vs. sequences in their immediate proximity (flanks). The analysis examined 31,319 instances across 11,034 genes. The X-axis measures the difference between the conservation scores of a pyknon instance and its immediate upstream or downstream flank. Positive values indicate that the pyknon instance is more conserved compared to its flanks, whereas negative values indicate the opposite. C: Boxplots showing the number of nucleotides (i.e., non-gaps) spanning the regions containing the 26,109 intronic instances of the 2,810 sequences of group_omni-TC_, group_omni-CGG_, and group_punct_.

Recall that rRNA/spacer pyknons have many genic copies. Indeed, the sense or antisense copies of the 1,723 pyknons across rRNAs, spacers, and the introns and exons of protein-coding genes span 2,436,551 nucleotides. The actual number of genomic positions covered by the 1,723 pyknons is even higher when one includes the pyknons’ *intergenic* exact copies.

For the pyknons (TC)_8_ and G(CGG)_5_ that lead group_omni-TC_ and group_omni-CGG_, respectively, we examined whether either is part of a wider motif encompassing additional nucleotides that perhaps differ from the “TC” and “CGG” cores. To this end, we enumerated the 180,955 human genome (T2T) copies of (TC)_8_ and G(CGG)_5_, respectively, flanked each copy with 20 nts on each side, and generated sequence logos^17^. **Supp. Figure S4** shows that the immediate neighborhoods of the genomic copies of these two pyknons involve a smaller, variable number of copies of the “TC” and “CGG” cores.

### Each rRNA and each spacer contains a unique set of pyknons that it shares with a unique set of mRNAs

We compared the pyknons found in each of the six rRNAs and four transcribed spacers of 45S (**Supp. Table S1C**) to determine whether they recur within these ten sequences. We carried out the analysis separately for the 875 sense and 880 antisense pyknons. **Tables 1A-B** show that any two of the ten reference sequences share no more than a handful of pyknons, independently of the pyknons’ orientation in the rRNAs or spacers.

We also compared the genes in whose mRNAs these pyknons are found and calculated the pairwise overlap of these gene groups. This calculation is informative for two reasons. First, even though a pyknon is constrained to be sense to at least one mRNA there is no guarantee that its remaining genomic copies exist in the mRNAs of other genes. Second, even though two rRNAs/spacers may share no pyknons, it is still possible that their respective pyknons co-occur in the mRNAs of the same gene(s). We additionally considered the relative orientation of pyknons in rRNAs/spacers and mRNAs. For example, a given pyknon can be sense to an rRNA/spacer and antisense to an mRNA. Examining pyknons that are *antisense* to mRNAs may seem paradoxical, considering that pyknons must have, by definition, one sense copy in an mRNA. However, note that many pyknons that are sense in one mRNA appear in antisense orientation in other mRNAs. To avoid multiple counting, we compared shared pyknons at the level of the genes, not mRNAs. **Table 2** summarizes our findings and includes several off-diagonal entries with high counts, the ETS1-ITS1, ETS1-28S, ITS1-ITS2, and ITS1-28S pairs. This crosstalk is expected and results from segments internal to these spacers/rRNAs that are duplicates or reverse complements of one another – see, for example, the red-colored region in the secondary structure of ITS1 and 28S in **Figure 1**.

### The sequences of group_omni-TC_, group_omni-CGG_, and group_punct_ appear in specific mRNA regions of specific genes

One or more sequences from group_omni-TC_, group_omni-CGG_, or group_punct_ are found in the mRNAs of 3,893 genes (**Supp. Table S2B**). We asked whether the 5’-UTR instances of these sequences favor specific gene groups. We then repeated the process for their CDS instances and, finally, their 3’-UTR instances. **Figure 4** and **Supp. Figures 5A-B** summarize an enrichment analysis^12^ at an FDR threshold of 1.0E-03. These Figures show that the rRNA/spacer motifs are prevalent in the mRNAs of nervous system genes and developmental genes that also have established associations with neuropathologies and genetic disorders.

**Figure 4.**
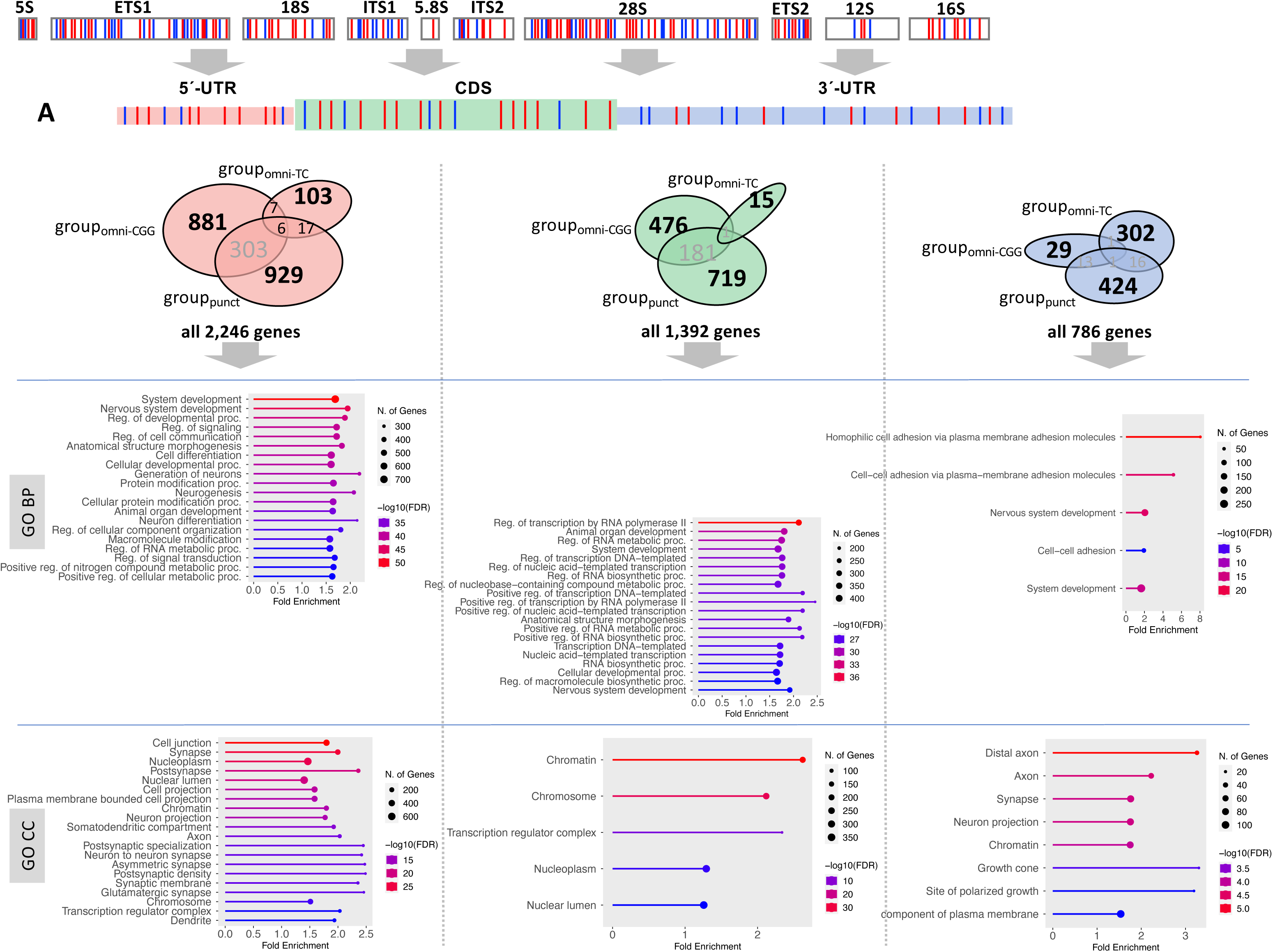
Enrichments of genes whose 5’-UTRs, CDSs, and 3’-UTRs share pyknons in sense (blue marks) or antisense (red marks) with rRNAs and 45S spacers. Shown are the enrichments for biological processes and cellular components.

We also asked whether the copies of the rRNA/spacer motifs we find in an mRNA region (5’-UTR, CDS, 3’-UTR) come from a specific group of sequences (group_omni-TC_, group_omni-CGG_, or group_punct_). **Supp. Figure S6** shows a division of labor among the nine possible combinations (3 mRNA regions x 3 groups of pyknons). Namely, each of group_omni-TC_, group_omni-CGG_, and group_punct_ contributes its sequences to a different mRNA region in largely non-overlapping sets of genes belonging to non-overlapping biological processes. Group_omni-TC_ and group_punct_ contributed the only exception: their sequences are in the 3’-UTRs of two sets of genes enriched in cell adhesion.

The entries of **Supp. Table S2A** and **Supp. Table S2B** also show that most of the copies of group_omni-TC_ sequences are in introns. Because group_omni-TC_ sequences are exclusively composed of pyrimidines, we examined whether they are an instance of the “polypyrimidine tract” that is typically located near and upstream of constitutive 3’ splice sites^18^. We found that only 1.9% of the group_omni-TC_ intronic copies are within 60 bases from the nearest downstream exon. This indicates that group_omni-TC_ sequences are unrelated to the canonical polypyrimidine signal.

### Only the genes whose pre-mRNAs pair up a sequence from group_omni-TC_ with a sequence from group_punct_ are linked to the nervous system and development

The entries of **Supp. Table S2B** allow three additional observations. First, far more genes contain sequences from only group_omni-TC_ or group_punct_ in their introns than their exons. Approximately one-half of the genes whose pre-mRNAs contain group_omni-TC_ sequences do *not* contain group_punct_ sequences, and *vice versa*. Enrichment analysis (FDR ≤ 1.0E-03) showed that the 2,941 genes that contain *exclusively* group_omni-TC_ sequences are involved in ion transport and metabolism (**Figure 5**). On the other hand, the 3,191 genes containing *exclusively* group_punct_ sequences are involved in diverse processes (FDR ≤ 1.0E-03).

**Figure 5.**
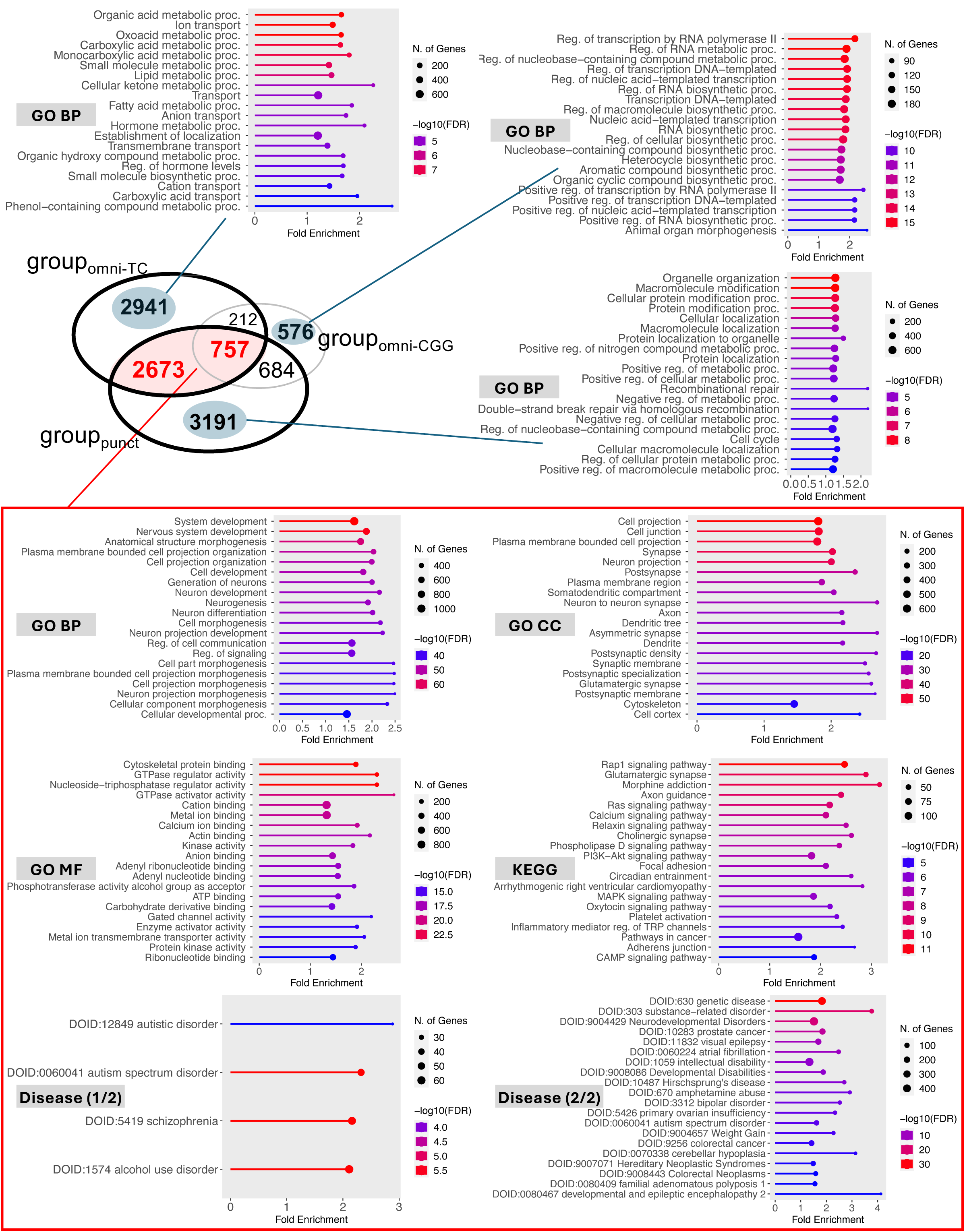
A: The three groups of sequences (group_omni-TC_, group_omni-CGG_, and group_punct_) appear in the pre-mRNAs of 11,034 genes. Most of these genes contain either a sequence from group_omni-TC_ or a sequence from group_omni-CGG_: only the genes that contain *at least one* sequence from each of these two groups are enriched in the nervous system and multiple neurological disorders. We generated the shown GO term enrichments with ShinyGO^12^ (FDR ≤ 1.0E-03). For clarity of presentation, not more than the top 20 terms of each list are shown.

Second, 3,430 genes simultaneously contain *at least one* sequence from each of group_omni-TC_ and group_punct_ in their pre-mRNAs. These 3,430 genes have strong (FDR ≤ 1.0E-03) multifaceted links to development and the nervous system. The enriched cellular component terms include axon, dendrite, and synapse (**Figure 5**). The enriched KEGG terms include signaling, axon, and synapse-related pathways. Moreover, there is a striking enrichment of disease terms, as determined by the two ontologies we examined, the Disease Alliance^19^ and the Disease Rat Genome Database^20^: 1,046 of the 3,430 genes are known risk loci for multiple neuropathologies and developmental disorders including autism, schizophrenia, bipolar disorder, epilepsy, and Hirschsprung’s disease.

Third, only 576 genes contain exclusively group_omni-CGG_ sequences in their introns (**Figure 5**). These genes are involved in transcription and RNA biogenesis, indicating that the role of group_omni-CGG_ sequences differs from that of the other two groups. This finding represents a novel link between rRNAs that have roles in translation and specific genes that have roles in transcription.

### The enrichments of the group_omni-TC_ and group_omni-CGG_ sequences remain after accounting for pre-mRNA length

The pre-mRNAs of nervous system genes are typically long^21^. This raises the possibility that the many genomic copies of group_omni-TC_ and group_omni-CGG_ sequences we find in these genes result by chance. To account for pre-mRNA length, we repeated the enrichment analysis of the genes containing group_omni-TC_ and group_omni-CGG_ sequences by including a gene’s genomic span as an independent covariate. We found that more than 99% of the significantly enriched GO biological process terms (FDR ≤ 1.0E-03, fold enrichment ≥ 1.5) of **Figure 5** remain statistically significant after accounting for length: 474/475 for group_omni-TC_ and 694/694 for group_omni-CGG_. Thus, the placement of these motifs in the pre-mRNAs of human genes is non-random.

### Each rRNA/spacer is associated with a different set of human pathologies

**Table 1** shows that each rRNA and spacer contains a unique set of pyknons. Moreover, **Figure 5** highlights the importance of the co-occurrence in pre-mRNAs of sequences from group_omni-TC_ and group_punct_. Given the genomic omnipresence of group_omni-TC_ sequences, we reasoned that it must be the group_punct_ sequences that establish the associations of rRNAs/spacers with different human pathologies. To test this, we identified the subset of group_punct_ sequences that are present in each rRNA and spacer (**Supp. Table S1C-D**), then identified which among the 3,430 genes of **Figure 5** pair up the subset with group_omni-TC_ sequences in their pre-mRNAs. We found strong associations (FDR ≤ 1.0E-03) between pathologies and the genes sharing group_punct_ sequences with the following rRNAs/spacers:

- 5S rRNA: The genes are associated with autism spectrum disorder (ASD).
- 28S rRNA: The genes are associated with substance-related disorders, neurodevelopmental disorders, epilepsy, bipolar disorder, ASD, atrial fibrillation, primary ovarian insufficiency, and prostate cancer.
- ETS1: The genes are associated with substance-related disorders, developmental disabilities, neurodevelopmental disorders, thrombocytosis, atrial fibrillation, and ASD.
- ITS1: The genes are associated with substance abuse, developmental disabilities, neurodevelopmental disorders, epilepsy, bipolar disorder, ASD, and prostate cancer.
- ITS2: The genes are associated with substance-related disorders, neurodevelopmental disorders, intellectual disability, epilepsy, and atrial fibrillation.
- ETS2: The genes are associated with substance abuse, neurodevelopmental disorders, Hirschsprung’s disease, thrombocytosis, epilepsy, atrial fibrillation, ASD, and colorectal cancer.

**Table 1.** Shared pyknons. The second row and second column show how many pyknons are sense (Table 1A) or antisense (Table 1B) in the six rRNA and four 45S spacers. The remaining rows and columns show how many of the pyknons of a given orientation are shared by any two of the ten regions. Note how the pyknons are unique to each rRNA and spacer.

| 1A |  | 5S | ETS1 | 18S | ITS1 | 5.8S | ITS2 | 28S | ETS2 | 12S | 16S |
| --- | --- | --- | --- | --- | --- | --- | --- | --- | --- | --- | --- |
|  |  | 6 | 250 | 34 | 121 | 2 | 102 | 333 | 25 | 2 | 10 |
| 5S | 6 | 6 | 0 | 0 | 0 | 0 | 0 | 0 | 0 | 0 | 0 |
| ETS1 | 250 |  | 250 | 0 | 0 | 0 | 0 | 5 | 2 | 0 | 0 |
| 18S | 34 |  |  | 34 | 0 | 0 | 0 | 0 | 0 | 0 | 0 |
| ITS1 | 121 |  |  |  | 121 | 0 | 2 | 0 | 0 | 0 | 0 |
| 5.8S | 2 |  |  |  |  | 2 | 0 | 0 | 0 | 0 | 0 |
| ITS2 | 102 |  |  |  |  |  | 102 | 0 | 0 | 0 | 0 |
| 28S | 333 |  |  |  |  |  |  | 333 | 3 | 0 | 0 |
| ETS2 | 25 |  |  |  |  |  |  |  | 25 | 0 | 0 |
| 12S | 2 |  |  |  |  |  |  |  |  | 2 | 0 |
| 16S | 10 |  |  |  |  |  |  |  |  |  | 10 |

| 1B |  | 5S | ETS1 | 18S | ITS1 | 5.8S | ITS2 | 28S | ETS2 | 12S | 16S |
| --- | --- | --- | --- | --- | --- | --- | --- | --- | --- | --- | --- |
|  |  | 10 | 242 | 35 | 116 | 0 | 125 | 332 | 30 | 2 | 1 |
| 5S | 10 | 10 | 0 | 0 | 0 | 0 | 0 | 0 | 0 | 0 | 0 |
| ETS1 | 242 |  | 242 | 0 | 0 | 0 | 0 | 5 | 2 | 0 | 0 |
| 18S | 35 |  |  | 35 | 0 | 0 | 0 | 0 | 0 | 0 | 0 |
| ITS1 | 116 |  |  |  | 116 | 0 | 2 | 0 | 0 | 0 | 0 |
| 5.8S | 0 |  |  |  |  | 0 | 0 | 0 | 0 | 0 | 0 |
| ITS2 | 125 |  |  |  |  |  | 125 | 3 | 0 | 0 | 0 |
| 28S | 332 |  |  |  |  |  |  | 332 | 3 | 0 | 0 |
| ETS2 | 30 |  |  |  |  |  |  |  | 30 | 0 | 0 |
| 12S | 2 |  |  |  |  |  |  |  |  | 2 | 0 |
| 16S | 1 |  |  |  |  |  |  |  |  |  | 1 |

**Table 2.**
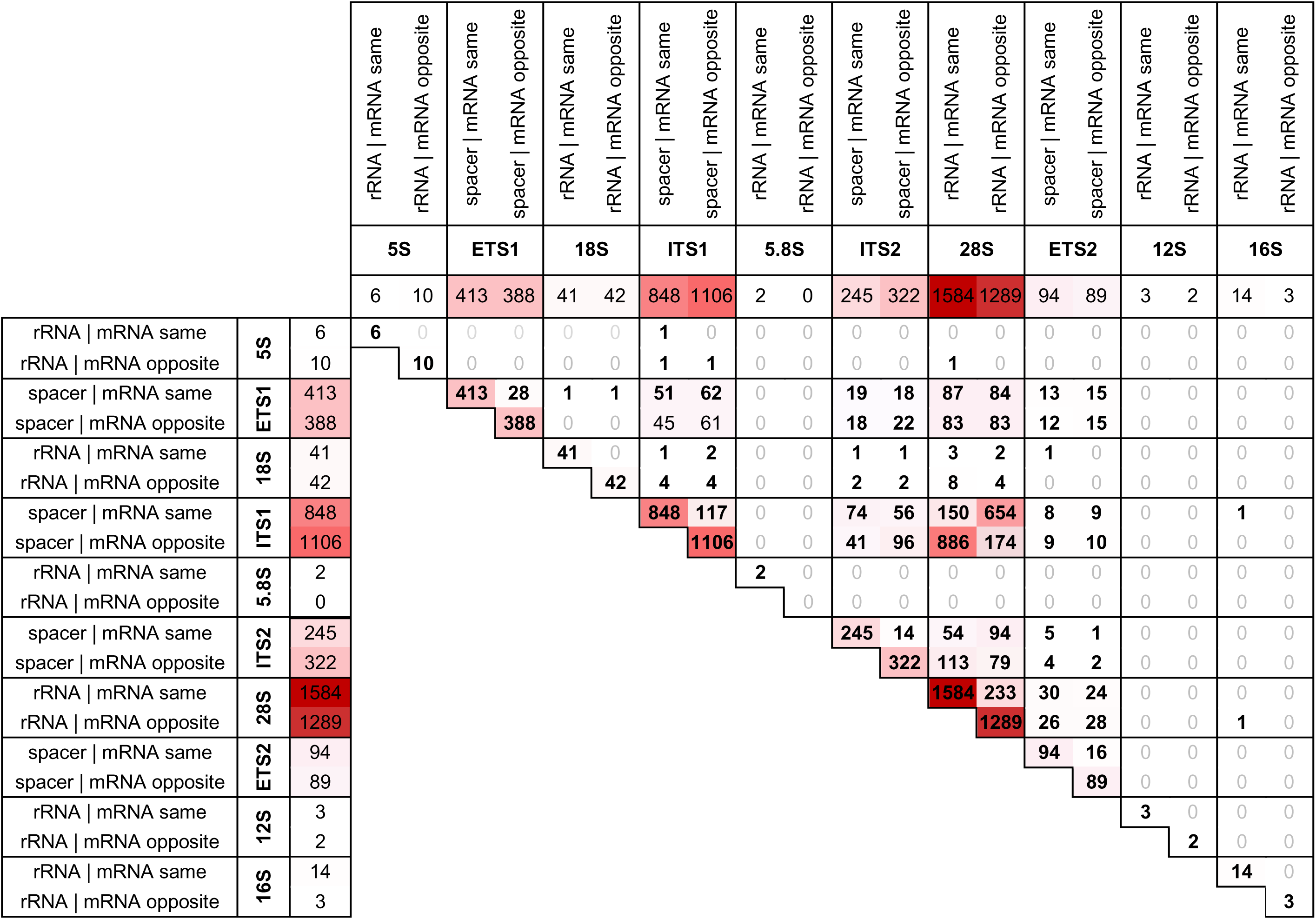
Number of genes linked to rRNAs and 45S spacers for all possible relative orientations. Off diagonal entries show the level of crosstalk among the various groups of genes. The Table accounts for 3,904 genes. Notation: “same” means same relative orientation; “opposite” means opposite relative orientation.

We found no notable associations for genes sharing group_punct_ sequences with the other four rRNAs/spacers. It is noteworthy that several disorders that are considered typically human (e.g., ASD, schizophrenia, bipolar disorder, Hirschsprung’s disease) are associated with the human-specific sequences of the transcribed spacers ETS1, ITS1, ITS2, and ETS2.

### Gene association with processes and disorders can depend on a pyknon’s orientation and be human-specific

The previous subsection highlighted distinct associations between pyknons found in rRNAs and 45S spacers and human pathologies. We examined this further with the help of the pyknon CGGGCGTGGTGGTGGG from ETS2 that belongs to group_punct_. Among the genes whose pre-mRNAs contain group_omni-TC_ pyknons, we identified those containing CGGGCGTGGTGGTGGG in sense (551 genes), antisense (671 genes), or both orientations simultaneously (171 genes). **Supp. Figure S7** shows that the relative orientation of this pyknon matters: each of the three gene groups is enriched for different biological processes, cellular components, molecular functions, and disorders. This ETS2 pyknon is human-specific. Therefore, its associations with nervous system genes are also human-specific (**Supp. Figure S7**).

### Group_omni-CGG_ sequences are the contact points for rRNA-mRNA and mRNA-mRNA heteroduplexes

Given the recurrence of exact sense and antisense pyknon copies in rRNAs, spacers, and mRNAs, we hypothesized that pyknons serve as points of direct interactions *in trans* between rRNAs and transcribed spacers on the one hand and mRNAs on the other. We analyzed data from three cell lines that were generated by the “psoralen-crosslinked, ligated, and selected hybrids” method (SPLASH)^22^ to test the hypothesis. First, we identified chimeric reads where one segment unambiguously mapped to an mRNA, and the remainder unambiguously mapped to an rRNA, a transcribed spacer of 45S, or a *different* mRNA. We sub-selected among the chimeric reads those containing any of the 1,723 pyknons in sense orientation in the first segment of the chimera and antisense orientation in the second segment, or *vice versa*, and computed enrichments compared to chance.

Strikingly, most heteroduplexes between rRNAs/spacers and mRNAs contain exact sense/antisense sequences from group_omni-CGG_ in the two segments of the chimeric reads. Frequently, the group_omni-CGG_ sequences appear in sense in the rRNA/spacer segment and in antisense in the mRNA segment. These pairings occur 10-100x more frequently than expected by chance (FDR ≤ 0.05, **Supp. Table S3A**). In all cells and biological replicates that we analyzed, the mRNA segments that participate in these hybrids are located primarily in the 5’-UTR and CDS regions (**Supp. Table S3B**). This result is anticipated by **Figure 4** and **Supp. Figures 5** for the sequences of group_omni-CGG_. We found very few chimeras supporting sense-antisense base-pairing of sequences from group_omni-TC_ and group_punct_.

Similarly, the mRNA-mRNA heteroduplexes that couple distinct mRNAs contain an over-representation of sense/antisense pairs of sequences from group_omni-CGG_ (FDR ≤ 0.05, **Supp. Table S3C**). As in the case of rRNA-mRNA heteroduplexes, most of these couplings involve 5’-UTRs (**Supp. Table S3D**). The 5’-UTR of Reelin (RELN) participates in most chimeras, coupled with the 5’-UTRs and CDSs of other mRNAs.

### The rRNA/spacer pyknons are also in the binding sites of numerous RBPs

We also investigated the possibility that RBPs interact with RNAs at pyknon sites. We searched all 1,723 rRNA/spacer pyknons in the CLIP-seq data of the 113 RPBs cataloged by the ENCODE project^23^. We sub-selected sites supported by ≥ 10 Reads-per-Million (RPM) whose support was ≥ 2x higher than observed in the matching control (IgG-CLIP) experiment. 941 (54.6%) of the 1,723 pyknons are fully embedded, in sense or antisense, in the binding sites of all 113 analyzed RBPs in various combinations (**Supp. Table S4**). These binding sites are in rRNAs, mRNAs, or pre-mRNAs. We also examined whether the pyrimidine-rich sequences of group_omni-TC_ are in the binding sites of PTBP1, U2AF1, or U2AF2, considering that these RBPs have an affinity for pyrimidine-rich regions^24,25^. Despite the genomic omnipresence of group_omni-TC_ sequences, for the RPM and fold-enrichment thresholds mentioned above, we found a mere 157 PTBP1 binding sites containing pyknons from this group (129 in K562 and 28 in HepG2 cells), 5 U2AF2 binding sites, and none for U2AF1. More than half of these sites are located in the 45S cassette, with only a few remaining sites occurring in introns or UTRs. This independent finding concurs with the earlier-mentioned finding that the group_omni-TC_ sequences are unrelated to the canonical polypyrimidine tract despite their intronic over-representation.

### Endogenous abundant rRNA-derived sncRNAs carry copies of rRNA/spacer pyknons

Given the presence of pyknons in rRNA/mRNA heteroduplexes and the binding sites of RBPs, we hypothesized that they instantiate molecular decoying in complete analogy to what miRNAs^3,26–28^ and tRNA-derived fragments (tRFs)^29^ are known to do. For this to occur, we would expect to find rRNA/spacer pyknons in the sequences of endogenous sncRNAs.

We searched 29,135 datasets from NIH’s Sequence Read Archive (SRA), all having passed quality control. We found that 1,222 (70.9%) of the 1,723 rRNA/spacer pyknons are carried in sense or antisense by 48,428 distinct sncRNAs. These sncRNAs are supported by ≥ 1,000 reads in at least one of the analyzed datasets.

In many settings, rRNAs and spacers produce sncRNAs known as rRNA-derived fragments or rRFs^30,31^. Thus, we hypothesized that the sncRNAs carrying the pyknons are rRFs and searched the recently released MINRbase^32^, which catalogs rRFs across many healthy and disease tissues. We found that 32,803 (67.7%) of the 48,428 sncRNAs are derived from rRNAs (rRFs), confirming our hypothesis. Most of the remaining pyknon-carrying sncRNAs arise from mRNAs, pre-mRNAs, or lncRNAs.

### Risk variants for neuropathologies overlap the genomic copies of rRNA/spacer pyknons

Having shown that rRNA/spacer pyknons are in rRNA-mRNA and mRNA-mRNA heteroduplexes, RBP binding sites, and endogenous sncRNAs, we hypothesized that their genomic copies must overlap known risk polymorphisms for the human disorders that we discussed above. We presume here that a risk variant occurring at a pyknon-containing location affects the affinity of the molecular interactions involving at least some of the pyknon’s remaining genomic copies.

To investigate this possibility, we examined whether the rRNA/spacer pyknons’ genomic copies overlap any variants listed in the NHGRI/EBI GWAS Catalog^33^. We only considered variants with an associated p-val ≤ 1.0E-08. We found 131 polymorphisms that overlap the genomic copies of 82 rRNA/spacer pyknons (**Supp. Table S5**): fold-enrichment=41.95 and p-val<3.9e-12. Most polymorphisms (90 out of 131) are found in genomic copies of sequences from the group_omni-TC_. The variants we identified are linked to neurological disorders, including autism spectrum disorder, schizophrenia, bipolar disorder, and depression, in agreement with our hypothesis. We note that the associations between GWAS-determined risk variants for diseases show considerable concordance with our findings mentioned above that we derived simply by analyzing the primary human genomic sequence.

### The sharing of *organism-specific* motifs between rRNAs/spacers and nervous system and developmental genes is conserved across evolution

We hypothesized that the sharing of rRNA/spacer pyknons with genes from nervous system and developmental genes is not unique to humans. Additionally, since the 45S cassette accounts for most of the human pyknons (**Supp. Figures S1-S2**), we reasoned that if such sharing persists in other organisms, it should be evident when only examining the 45S sequence.

We selected three common model organisms for this analysis: mouse, fruit fly, and worm. Because the fruit fly and the worm have small genomes (140 and 100 Mbps, respectively), we reasoned that the counterpart pyknons comprise fewer than 16 nts. Using a Monte Carlo simulation, we determined the probability of a random 13-mer appearing by chance 30 or more times in the fruit fly or the worm to be < 1.0E-03. For simplicity, we used redundant pyknons in these calculations (see Supplemental Methods). We also removed pyknons containing 15 A’s or 15 T’s.

Based on their genomic counts, we divided each organism’s pyknons into two groups: the counterpart of the human (TC)_8_ group, and all others. Since we already showed (**Figure 2**) that the sequences of group_omni-CGG_ are a recent innovation of primates, we did not seek a counterpart of group_omni-CGG_ in the genomes of the three model organisms. To create the counterparts of the human group_omni-TC_ and group_punct_ sets, we added the respective pyknons’ reverse complement to each pyknon collection and removed duplicates.

The mouse counterparts to the human group_omni-TC_ and group_punct_ sets contain 24 and 172 sequences, respectively (**Supp. Table S6A**). The fruit fly counterparts contain 272 and 2,540 sequences, respectively (**Supp. Table S6E**). And the worm counterparts contain 34 and 268 sequences, respectively (**Supp. Table S6I**). There are 10,208 mouse genes, 6,802 fruit fly genes, and 6,561 worm genes containing sequences from *either* group_omni_ or group_punct_ in their pre-mRNAs (**Supp. Table S6B-C, S6F-G, S6J-K**).

In complete analogy to what we found in humans (**Figure 5**), in all three model organisms only those genes whose pre-mRNAs contain *at least* one pyknon from group_omni_ and *at least* one pyknon from group_punct_ are associated with nervous system and developmental genes, and neurological and genetic disorders. **Supp. Figure S8**, **Supp. Figure S9**, and **Supp. Figure S10** show the corresponding pathway enrichments associated for mouse, fruit fly, and the worm, respectively. In each case, the enriched terms mirror those of the human genome (**Supp. Table S6D, S6H, S6L**). In the worm, the three gene sets captured by the Venn diagram of **Supp. Figure S10** show modest pathway crosstalk; however, the enrichments and statistical significance of the intersection (genes containing at least one pyknon from each of group_omni_ and group_punct_) far outweigh this crosstalk.

Lastly, we examined whether a model organism’s 45S cassette pyknons are in the 45S cassette of the other three organisms. To ensure generality, we considered the sense and antisense instances of the pyknons in each organism’s 45S cassette: there are 2,754 human, 196 mouse, 2,812 fruit fly, and 302 worm such sequences. **Table 3** shows the results of the pairwise tests. Each organism’s rRNAs and 45S spacers share different sequences with the organism’s nervous system genes, reinforcing the observation that these linkages are organism-specific.

**Table 3.** Searching the pyknons found in the 45S cassette of the human, mouse, fruit fly, and worm genomes in sense or antisense in the 45S cassette of the other two genomes reveals organism-specific sequences.

|  |  | TARGET |  |  |  |
| --- | --- | --- | --- | --- | --- |
|  |  | *H. sapiens*<br>(45S cassette) | *M. musculus*<br>(45S cassette) | *D. melanogaster*<br>(45S cassette) | *C. elegans*<br>(45S cassette) |
| **S**<br>**O**<br>**U**<br>**R**<br>**C**<br>**E** | *H. sapiens*<br>pyknons in 45S<br>(16 nts) | **2,754** | 418 | 56 | 48 |
|  | *M. musculus*<br>pyknons in 45S<br>(16 nts) | 12 | **196** | 2 | 0 |
|  | *D. melanogaster*<br>pyknons in 45S<br>(13 nts) | 120 | 124 | **2,812** | 54 |
|  | *C. elegans*<br>pyknons in 45S<br>(13 nts) | 4 | 12 | 8 | **302** |

## DISCUSSION

We focused on and analyzed a specific type of genome-wide-conserved motifs, the pyknons, and their presence in the six rRNAs (5S, 5.8S, 12S, 16S, 18S, 28S) and the four transcribed spacers of 45S (ETS1, ITS1, ITS2, ETS2). We showed that most of these sequences are densely packed with organism-specific recurring genomic motifs that they share with the pre-mRNAs of 11,034 human genes. Unlike previous work that sought similarities between full-length rRNAs and genes^34,35^ or used variable-length k-mers^36,37^ but studied only the expansion segments^38^ of 18S and 28S, we examined the four nuclear rRNAs, the four transcribed spacers of 45S, the two mitochondrial rRNAs, intronic sequences, possible links to disease, and conservation across evolution. Moreover, we coupled our analyses to experimentally-determined RNA-RNA interactions, RBP assays, transcriptomic data from tens of thousands of public datasets, and risk variants from genome-wide association studies. We uncovered a specific combination of rRNA and spacer motifs that occurs primarily in the pre-mRNAs of nervous system and developmental genes, including many risk genes for several complex genetic and neurological disorders. We also showed that this sharing of motifs between rRNAs/spacers and human nervous system genes is not unique to humans. It is also present in three model organisms (mouse, fruit fly, worm), with each organism using its own set of motifs. The persistence of this observation in four organisms separated by 600 million years of evolution suggests that it represents an important component of each organism’s development. The findings implicate for the first time the sequences of rRNAs and 45S spacers in the nervous system, developmental processes, and human pathologies through primate-specific motifs shared by rRNAs/spacers and specific genes. The motifs’ non-random genomic placement highlights yet again the importance of genomic architecture. Moreover, the data indicates that these motifs are central to rRNA/mRNA, mRNA/mRNA, RBP/rRNA, and RBP/mRNA interactions. Our study also suggests a role for the rRFs, a recently reported class of sncRNAs that are derived from rRNAs and spacers^31,32,39^: pyknon-carrying rRFs can modulate these interactions and be disrupted by risk variants at the genomic copies of pyknons. Below we will discuss the implications of these findings for the genetic dimension of neurological disorders and the creation of novel diagnostics and therapies.

While we only examined the pyknons in rRNAs and the spacers of 45S, our approach generalizes to lncRNAs that contain organism-specific pyknons. Based on initial evidence^5^, these lncRNAs could also use decoying to control gene groups within and beyond the nervous system through shared pyknon sequences. We expect that going beyond rRNAs and the spacers of 45S and extending the model to include lncRNAs will provide additional insights into the primate-specific dimension of other human conditions and diseases.

Each of the ten human rRNAs and spacers we analyzed contains a unique set of pyknons (**Table 1**), in sense or antisense (**Supp. Table S1A-B**). With the exception of 12S and 16S, the rRNAs and spacers are unusually rich in pyknons (**Supp. Figures S1-S2**). **Figure 1** shows the secondary structures for two rRNAs and two spacers with the pyknon-covered portions highlighted. The 1,723 pyknons found in rRNAs/spacers form three groups (**Supp. Table S1C**): the pyknon (TC)_8_ and its variants; the pyknon G(CGG)_5_ and its variants; and all remaining pyknons. By adding the pyknons’ reverse complements we formed and studied three sets of sequences, group_omni-TC_, group_omni-CGG_, and group_punct_ (**Supp. Table S1E**). **Figure 1** and **Supp. Figures S1-S2** show that not all pyknon-covered regions correspond to expansion segments^38^, nor are all expansion segments covered by pyknons.

We analyzed the instances in 28S of (TC)_8_ and G(CGG)_5_ and showed that they represent very recent insertions into the human genome (**Figure 2**). Indeed, most of their copies are absent from *N. leucogenys,* suggesting that the insertions occurred sometime in the last 15-20 million years. The recency of these insertions extends to *all* the pyknons we find in rRNAs/spacers and their numerous copies in the exons and introns of protein-coding genes (**Figure 3A-B** and **Supp. Figure S3A**). Indeed, the average span of the intronic and 3’-UTR copies of these human pyknons decreases to zero in their non-primate orthologs (**Figure 3C** and **Supp. Figure S3B**).

We particularly note the multiple instances of the trinucleotide CGG and its variants in 28S (**Figure 2**) and the 5’-UTRs and CDSs of genes (**Supp. Table S2B** and **Supp. Figure S3A**). CGG runs were previously discussed^40,41^ in the context of neuropathologies that are typically human^40,42–45^. **Figure 2** shows that the copies of G(CGG)_5_ in 28S occur only in primates. The only exception is the 28S rRNA of the odontocete *T. truncatus* (bottlenose dolphin), a non-primate mammal known to develop Alzheimer’s disease-like pathology^46^; it contains one copy of G(CGG)_5_. The other copies of G(CGG)_5_ are in the human ITS1 (in antisense – see **Figure 1**), the latter’s sequence being unique to primates that are evolutionarily close to humans (data not shown).

**Figures 2-3** and **Supp. Table S1C** capture an important corollary of our study. The sense and antisense copies of the 1,723 pyknons in rRNAs and spacers occur at tens of thousands of locations in the introns and exons of specific genes. Our finding that most of these genic copies are unique to primates suggests that these insertions occurred in recent evolutionary times and were coordinated across many sites and chromosomes.

Even if we confine our analysis to only the exonic instances of the rRNA/spacer pyknons, there is a clear “division of labor” that is evidenced by **Table 2**, **Figure 4**, and **Supp. Figure S5.** According to **Table 2**, the pyknons link each rRNA and spacer to the mRNAs of its own set of genes. The length of an rRNA/spacer and the number of genes with whose mRNAs it shares pyknons are not correlated. For example, the short (1,077 nts) internal transcribed spacer ITS1 is associated with 10x more genes than the longer 16S (1,559 nts) and 18S (1,869 nts). As another example, while the 28S rRNA is only 3x longer than the 18S rRNA, it is associated with ∼30x more genes.

Looking at enrichments, **Supp. Figure S5** shows that each of the nine possible combinations of {group_omni-TC_, group_omni-CGG_, group_punct_} x {5’-UTR, CDS, 3’-UTR} corresponds to genes that are associated with different biological processes. Even when we combine together all the genes whose 5’-UTRs, CDSs, and 3’-UTRs contain *any* of the sequences in group_omni-TC_, group_omni-CGG_, group_punct_, we arrive at a similar observation (**Figure 4**).

These genome-driven analyses led to three novel observations that are unique to primates:

- Genes whose *exons* contain group_omni-TC_ sequences are linked to RNA splicing even though the *intronic* copies of group_omni-TC_ sequences are unrelated to the polypyrimidine tract/splicing (**Supp. Figure 6**).
- Genes whose exons contain group_omni-CGG_ sequences are involved in DNA transcription by RPOL II (**Supp. Figure 6**).
- Genes whose 5’-UTRs or 3’-UTRs contain group_omni-TC_, group_omni-CGG_, or group_punct_ sequences are linked to multiple genetic disorders and neuropathologies (**Supp. Figure 5**).

The first two observations link transcription and splicing on the one hand with rRNAs and translation on the other. The third observation links rRNAs and translation with multiple human disorders. The primate-specificity of these linkages result from the fact that the sequences of group_omni-TC_, group_omni-CGG_, and group_punct_ exist only in the rRNAs and spacers of primates.

Another key finding of our study is the evident “cooperation” of the sequences in group_omni-TC_ and group_punct_. The analysis shows that a necessary and sufficient condition for a gene to be linked to development and the nervous system is to contain *at least one* pyknon from each of these two groups (**Figure 5**). The cooperating nature of pyknons from these two groups is elaborate. When we pair group_omni-TC_ pyknons with only the sequences of group_punct_ found in a single rRNA/spacer, we end up with a *different* set of genes each time, and the set is linked to a different group of diseases/conditions. This suggests a “division of labor” and reinforces our choice of the characterization “punctate” for the pyknons of this group. This division was presaged by the results shown in **Table 2**.

To reiterate, there is a solid primate-specific dimension to the linkages between genes and the sequences in group_punct_. A human-specific pyknon, CGGGCGTGGTGGTGGG from ETS2, helped highlight this. Genes that pair up group_omni-TC_ sequences with this pyknon in sense orientation are enriched for different biological processes, cellular components, molecular functions, and diseases than genes that pair up group_omni-TC_ sequences with the pyknon in antisense (**Supp. Figure S6**).

The recurrence of exact sense and antisense pyknon copies from rRNAs/spacers in numerous genes raised the possibility that pyknons are involved in long-range interactions. Evidence already exists for interactions based on direct sense/antisense coupling^47,48^, as well as for interactions driven by decoying sncRNAs^5,26–29,49^. We wanted to examine whether any such interactions involve rRNA/spacer pyknons.

- direct-coupling: We analyzed psoralen-crosslinked chimeras from several cell lines and replicates. We found multiple chimeras where one segment unambiguously contained a rRNA-derived sequence containing a pyknon and the other an mRNA fragment containing the pyknon’s reverse complement (**Supp. Table S3**). We also found multiple chimeras where the two segments corresponded to different coupled mRNAs (**Supp. Table S3**). Most of the pyknons participating in these chimeras were from group_omni-CGG,_ and most mRNA fragments were from 5’-UTRs and CDSs.
- decoying interactions (1): We searched for instances of the 1,723 pyknons in the binding sites of RBPs. We found that in various combinations, 79.9% of the pyknons from rRNAs and spacers are in the binding sites of all 113 RBPs reported by the ENCODE project. One possible use of such pyknon embeddings could involve rRNAs/spacers and mRNAs containing pyknons in the same orientation and competing for binding to the same RBP^50^.
- decoying interactions (2): We searched 29,126 public datasets from NIH’s SRA repository for sncRNAs containing any of the 1,723 pyknons from rRNAs/spacers in either sense or antisense. We found 1,222 (70.9%) of these pyknons carried by 48,428 sncRNAs, each having an abundance of 1,000 reads or more in at least one dataset. 30.8% of these sncRNAs are produced by lncRNAs and the exons and introns of mRNAs^47,48,51,52^. This suggests the possibility of molecular decoying of rRNA/mRNA and RBP/RNA interactions by rRNA-derived pyknon-containing sncRNAs.

Many of the pyknons we find in the rRNAs and spacers also overlap previously reported clinically-significant variants, reinforcing the importance of these multifaceted *cis* and *trans* relationships. Searching the GWAS Catalog showed that 131 risk variants with associations to many of the same disorders and conditions that arise from our genome-only analyses overlap genomic copies of 82 of the 1,723 pyknons (p-val=3.9e-12) (**Supp. Table S5**). This agreement further highlights two points. The first point is a pervasive observation in our presentation, namely, that there is a strong connection between the architecture of the genome, the transcriptome, and known disorders. The second point is that the presence of the variants within pyknon copies highlights the pyknons’ functional relevance and goes beyond a simple association with disorders.

This second point is corroborated by the recurrent presence of these motifs in a) sncRNAs from thousands of independently generated samples from different tissues and donors, b) RNA-RNA heteroduplexes, and c) RBP binding sites.

The recurrent presence of these motifs in sncRNAs indicates consistency and suggests a purpose. However, our understanding of this purpose is currently limited. The evidence suggests a model where sncRNAs use “sponging” or “decoying” to regulate the formation of RNA-RNA heteroduplexes or the binding of RBPs through shared motifs. For example, sncRNAs could affect the direct coupling between RNAs that contain pyknons in sense/antisense orientations and would normally facilitate the direct recruitment of ribosomes on mRNAs to promote their translation. This mode is supported by our finding of the motifs at the rRNA-mRNA contacts in psoralen crosslinking data^22^. Recent work by Barna and colleagues suggests that a similar mechanism is operating in human neuronal cells^53^. Similarly, sncRNAs could affect the binding of RBPs to sites of mRNAs/pre-mRNAs that contain pyknon copies, by extension influencing RNA stabilization and degradation, pre-mRNA splicing, helicase activity, localization, transport, and polyadenylation^50,54,55^. These possibilities suggest that the intronic and exonic copies of pyknons are functional pre-mRNA/mRNA features.

The “rRNA/spacer–genome architecture–nervous system” axis is not unique to humans. Indeed it also exists in mice, fruit flies, and worms, and thus, spans 600 million years of evolution. In all cases, the axis leverages organism-specific motifs that are shared between the organism’s rRNAs and spacers and the organism’s nervous system and developmental genes. The 45S cassettes of mice, fruit flies, and worms are densely packed with pyknons that can be separated into two sets, the counterparts of the human group_omni_ and group_punct_ sets. The sequences comprising these counterpart sets are organism-specific (**Table 3**). And just like in humans, the model organism genes whose pre-mRNAs contain at least one pyknon from each of the organism’s group_omni_ and group_punct_, and only those genes, are involved in developmental and nervous system processes, and linked to genetic disorders and neuropathologies (**Supp. Figures S7-S8**). These findings strengthen the observation that the placement of these genome-wide, organism-specific motifs across the genomes of these three organisms is not accidental.

A particularly intriguing aspect of our findings is that the pyknons described herein establish a novel, tangible connection between rRNAs/spacers and mRNAs that have been implicated in diseases characterized by aberrant ribosome biosynthesis or deficient ribosomal function collectively called ribosomopathies^56^. A characteristic aspect of these diseases is that although defective ribosome biosynthesis and function is ubiquitous, it appears to be particularly deleterious in specific tissues and cell types, often leading to neurodevelopmental impairment, craniofacial deformities, bone marrow and cardiac defects, and other issues^56–58^. Several genes with a role in ribosomopathies appear in the three groups of pyknons and the intersection of group_punct_ with either of the other two, such as POLR1C and POLR1D (Treacher Collins syndrome)^59^, POLR1A^60^, UTP4^61^, DNAJC21^62^, TRPS1, many ribosomal protein genes, DDX51, DDX17, and DDX47 RNA helicases involved in ribosome biosynthesis. Moreover, although insofar not directly linked with ribosomopathies, many other factors involved in mRNA translation and its regulation are present in our three groups, such as eIF2alpha kinases HRI, PKR, and PERK and translation initiation and elongation factors. Mitochondrial ribosomal protein genes implicated in various cancers are also found in those lists; it is worth noting that ribosomopathies often show a propensity for cancer^56,58,63^. Finally, emerging evidence connects many additional neurological disorders with mRNA translation aberrations, such as autism and Rett syndrome^64,65^, which also appeared in our GO analysis. The sharing of pyknons between rRNA and mRNAs involved in ribosomopathies constitutes a novel molecular link that could provide a new basis for exploring the poorly understood pathogenetic mechanisms of many ribosomopathies.

We stress the existence of associations between rRNAs/spacers and genes that are known risk loci for autism. These findings make a strong argument for the genetic basis of this condition. Moreover, in the cases of autism, schizophrenia, and bipolar disorder – all ailments that typically affect humans – the absence of syntenic conservation of human pyknons beyond primates suggests a new vantage point from which to study the mechanisms that underlie these disorders. If supported by future findings, these associations would help create much-sought molecular diagnostics. A related finding is the presence of the rRNA/spacer pyknons in many genes linked to substance-related disorders. Just like autism, the finding suggests a genetic basis for substance abuse^66–68^. We posit that polymorphisms, duplications, micro-insertions, or micro-deletions at the sites of these pyknons, or their immediate neighborhood, may underlie the two conditions.

In closing, we reiterate our inkling that the linkages between genome architecture, biological processes, and human disorders that we uncovered are not unique to rRNAs/spacers. We expect that different lncRNAs will contain different sets pyknons^5,7^ in different orientations, creating intragenomic linkages of specific lncRNAs to specific groups of genes. The many human pyknons, the high numbers of their genomic copies, the many annotated protein-coding genes, and the many long noncoding RNAs suggest numerous potential linkages that are currently uncharacterized.

## Author Contributions

IR designed and supervised the study. IR, SN, PL, IN, BD, EL, and AV generated and analyzed the data. IR wrote the paper with assistance from SN, PL, AV, and EL.

## SUPPLEMENTAL FIGURE CAPTIONS

**Supp. Figure S1.** Regions of the six rRNAs and four spacers of 45S containing pyknons in sense orientation. Overlapping pyknons cover the segments shown in lowercase.

**Supp. Figure S2.** Regions of the six rRNAs and four spacers of 45S containing pyknons in antisense orientation. Overlapping pyknons cover the segments shown in lowercase.

**Supp. Figure S3.** Conservation of group_omni-CGG_ instances in 5’-UTRs and CDSs. A: Multiple sequence alignments of syntenic regions from four representative genes. The human sequence contains an instance from group_omni-GCC_ (corresponding nucleotides are highlighted in green; partial CGG copies are colored yellow). The examples show a lack of conservation in organisms that are phylogenetically distant from humans. B: Boxplots show the width of the region that corresponds to the 5,210 instances in 5’-UTRs, CDSs, and 3’-UTRs (see **Figure 2**) of the 2,810 sequences from group_omni-TC_, group_omni-GCC_, and group_punct_.

**Supp. Figure S4.** Logos of 20-nt-long regions that immediately flank the G(CGG)_8_ and (TC)_8_ pyknons across the entire genome.

**Supp. Figure S5.** Enrichments of genes whose 5’-UTRs, CDSs, and 3’-UTRs share pyknons in sense (blue marks) or antisense (red marks) with rRNAs or 45S spacers. Enrichments for molecular functions, KEGG pathways, and diseases.

**Supp. Figure S6.** The pyknons in rRNAs and spacers form three groups (group_omni-TC_, group_omni-CGG_, and group_punct_). The sequences in these groups have specific distributions in the 5’-UTRs, CDSs, and 3’-UTRs and are found in specific groups of genes. We generated the shown GO term enrichments with ShinyGO^12^ (FDR ≤ 1.0E-03). For clarity of presentation, not more than the top 10 terms of each list are shown. Blue and red marks represent pyknons in sense and antisense orientation, respectively.

**Supp. Figure S7.** Pyknon orientation matters. We formed two groups of genes: those whose pre-mRNAs contain both the ETS2 pyknon CGGGCGTGGTGGTGGG and at least one group_omni-TC_ sequence in sense; and those whose pre-mRNAs contain the ETS2 pyknon CGGGCGTGGTGGTGGG in antisense and at least one group_omni-TC_ sequence in sense. We determined enrichments for each of the three subsets of genes defined by the Venn diagram (FDR ≤ 1.0E-03). As can be seen, each combination of pyknon orientation is linked to a different set of biological processes, cellular components, molecular functions, and diseases.

**Supp. Figure S8.** The sequences of the mouse group_omni_ and group_punct_ exhibit the same properties as their human counterparts: only the mouse genes that contain *at least one* sequence from each of these two groups are enriched in nervous system and developmental genes, and multiple genetic and neurological disorders (FDR ≤ 1.0E-03). For clarity of presentation, only the top 20 terms of each list are shown. Blue and red marks represent pyknons in sense and antisense orientation, respectively.

**Supp. Figure S9.** The sequences of the fruit-fly’s group_omni_ and group_punct_ exhibit the same properties as their human and mouse counterparts: only the fruit fly genes that contain *at least one* sequence from each of these two groups are enriched in nervous system and developmental genes, and multiple genetic and neurological disorders (FDR ≤ 1.0E-03). For clarity of presentation, only the top 20 terms of each list are shown. Blue and red marks represent pyknons in sense and antisense orientation, respectively.

**Supp. Figure S10.** The sequences of the worm’s group_omni_ and group_punct_ exhibit the same properties as their human, mouse, and fruit-fly counterparts: the worm genes that contain *at least one* sequence from each of these two groups are enriched in nervous system and developmental genes (FDR ≤ 1.0E-03). For clarity of presentation, only the top 20 terms of each list are shown. Blue and red marks represent pyknons in sense and antisense orientation, respectively.

