## Supplemental Methods for "All ribosomal RNAs and 45S spacers from humans to worms are packed with organism-specific motifs whose other copies are found predominantly in numerous nervous system genes including many associated with human disorders"

**Exhaustive enumeration of genomic k-mers (k=16)**

In our previous work^1, 2^ we used a pattern discovery method^3^ to identify variable *maximal*-length motifs of length ≥ 16 nts with ≥ 30 identical copies in the intergenic/intronic regions of human and mouse. While we discovered motifs longer than 16 nts, ~96% of the pyknons in the final collection were exactly 16 nts long in both organisms. Considering this and the high computational burden of discovering maximal-length patterns, we simplified this step by identifying 16-mers with ≥ 30 identical copies in the T2T genome^4^. We also modified the requirement and now allow a pyknon to have ≥ 30 *genomic* instances instead, at least one of which is in an mRNA. This modification is meant to accommodate the possibility that some pyknons may have their genomic instances primarily, if not exclusively, *inside* mRNAs. In these analyses, we did not include the Y chromosome of the T2T genome to avoid potential bias in favor of male-specific motifs.

**Statistical significance of the enumerated 16-mers**

By fixing the length to k=16 and requiring a minimum number of genomic copies, we can ensure the statistical significance of the pyknons. Indeed, using a Monte Carlo simulation, we determined the distribution of 16-mers in randomly shuffled instances of the T2T genome assembly. Specifically, we rearranged the forward strand, then generated the reverse complement of the shuffled instance, and enumerated all distinct 16-mers in the resulting two-stranded genome. We repeated this process 100 times, each creating a histogram of the frequencies of the 16-mers in the respective shuffled instance of the genome.

**Identification of pyknons among the genomic 16-mers**

Pyknons are constrained to have at least one of their genomic copies in an mRNA. To this end, we first filtered the 16-mers with ≥ 30 identical copies in the T2T genome and kept only those that are also present in at least one of the 96,441 mRNAs (22,511 protein-coding genes) of Rel. 105 of ENSEMBL. The 16-mers that survived this step formed a list of “*candidate pyknons*.” Using the approach outlined in our original study^1^, we identified the final list of pyknons by removing *redundant* 16-mers among these candidates. Two fixed-length pyknons are redundant if they have the same genomic counts and co-occur at the same genomic locations. For example, suppose the 17-mer CCCATACCACGGGGATT is conserved genome-wide. In that case, it will give rise to two 16-mers, CCCATACCACGGGGAT and CCATACCACGGGGATT, with the same counts whose overlap is CCATACCACGGGGAT: keeping one of the two 16-mers suffices. Briefly, the redundancy removal proceeds as follows. We begin by marking all nt positions in all ENSEMBL mRNAs as “not covered” by a pyknon. We sort all candidate pyknons in order of decreasing number of their genomic copies. Candidate pyknons with the same count are sorted lexicographically. For each candidate, in turn, we seek instances of it in the mRNA sequences. If the candidate is found in an mRNA and all positions of its match are labeled “not covered,” then the candidate “claims” these positions and changes their labels to “covered.” If we can place the candidate pyknon more than once in the current mRNA, we do so before examining the next mRNA. Each candidate that successfully claims one or more spots in one or more mRNAs during this traversal is reported as a “pyknon”; otherwise, it is discarded.

**Reference rRNAs**

For this analysis, we used the following human genome reference sequences from GenBank: RNA45SN1 (accession number NR_145819.1) as the reference entry for 45S; RNA5S12 (accession number NR_023374.1) for 5S; MT-RNR1 (accession number NR_137294.1) for 12S; and MT-RNR2 (accession number NR_137295.1) for 16S. These reference sequences have the following lengths: RNA45SN1 - 13,351 nts; RNA5S12 - 121 nts; MT-RNR1 - 954 nts; MT-RNR2 - 1,559 nts, respectively. Within 45S, we distinguished among the 5´ ETS or ETS1, 18S rRNA, ITS1, 5.8S rRNA, ITS2, 28S rRNA, and 3´ ETS or ETS2. We also used the following 19 non-human 28S rRNA reference sequences: NR_000055.1 (*C. elegans*), NR_003279.1 (*M. musculus*), NR_046246.2 (*R. norvegicus*), NR_133562.1 (*D. melanogaster*), XR_002740182.2 (*F. catus*), XR_002805825.1 (*E. caballus*), XR_003510171.1 (*B. indicus x B. taurus*), XR_003725683.1 (*M. mulatta*), XR_003887306.1 (*T. rubripes*), XR_004030810.1 (*N. leucogenys*), XR_004223802.1 (*X. tropicalis*), XR_004524865.1 (*T. truncatus*), XR_005382296.1 (*C. lupus familiaris*), XR_005582245.1 (*S. boliviensis boliviensis*), XR_005647714.1 (*P. giganteus*), XR_006938338.1 (*G. gallus*), XR_007912736.1 (*O. cuniculus*), XR_008546126.1 (*P. troglodytes*), and XR_008676494.1 (*G. gorilla*). Lastly, we used the following five non-human precursor 48S rRNA reference sequences: KX061886.1 (*P. troglodytes*), KX061888.1 (*P. abelii*), KX061890.1 (*M. mulatta*), KX061887.1 (*G. gorilla*), and KX061889.1 (N. leucogenys). The reference sequences for 45S for the *M. musculus*, *D. melanogaster*, and *C. elegans* genomes are NR_046233.2, NR_133558.1, and X03680.1, respectively.

**Statistical significance of pyknons found in rRNAs/spacers, mRNAs, and pre-mRNAs**

For each T2T pyknon found in the ten reference sequences (six rRNAs and four spacers), we determined whether it is enriched in each of 5S, 45S, 12S, 16S, 5´ untranslated regions (5´-UTRs), coding sequences (CDSs), 3´ untranslated regions (3´-UTRs), mRNAs, introns, and pre-mRNAs, respectively. To this end, we compared a pyknon’s observed sense or antisense counts in each region to its expected counts and computed fold enrichments and p-values using a binomial test.

**Enrichment analysis of gene sets**

We used ShinyGO^5^ to find enriched GO terms, KEGG pathways, and The Alliance of Genome Resources diseases in the input gene lists (FDR ≤ 0.001; fold enrichment ≥ 1.5). We used REVIGO^6^ to visualize the enrichments of GO terms. To test a pathway enrichment in a given gene list adjusted for genes’ lengths, we adapted a previously proposed approach^7^. We considered a logistic regression model, where the indicator of pathway belonging is a dependent variable, and the indicator of gene list belonging and gene length are independent variables. We computed a gene’s length as averaged pre-mRNA lengths across all transcripts of the gene. We fit the regression model and get p-values for independent variables using the *statsmodels* v0.14.0 Python module.

**Expanding the pyknon groups**

We split the set of the pyknons we find in rRNAs and spacers into two subsets: the first is the pyknons we want to expand, and the second all remaining pyknons of the set. We iteratively augmented the first set by allowing each member to “attract” pyknons from the second set if the two have a Levenshtein distance ≤ 1. The process proceeds iteratively, moving pyknons to the set being expanded, and terminates when the first set can attract no more pyknons.

**Searching for syntenic sequences containing pyknons and quantifying sequence conservation**

For each pyknon instance of interest from the set of 175 pyknons (*expanded* G(CGG)_5_ and (TC)_8_ groups) within 5´-UTRs, CDSs, 3´-UTRs, or introns of a gene, we determined its syntenic region in each of the above-mentioned 19 non-human genomes as follows. First, we extended the pyknon instance both upstream and downstream to include any additional copies of the 175 pyknons located ≤ 20 nts from each other. Second, we flanked the extended sequence on the genome with 100 nts on each side (we did not analyze the few pyknon instances that overlap exon-exon junctions). Third, we sought the resulting sequence (pyknons+flanks) in the genome of each organism in the proximity (±100 kb) of the orthologous gene. We performed the search using the GLSEARCH v36.3.8i^8^ with the default E-value threshold and ≥ 75% nucleotide identity threshold; we considered only the best alignment (if any existed). We multiply-aligned the resulting sequences using clustal-omega^9^. For each position on the multiple alignment, we calculated a conservation score using the formula^10^: score = (1 - scaled Shannon’s entropy) x (1 - fraction of gaps). We averaged the conservation scores in the window of positions covered by the pyknon instances in humans and in the flanking alignment portions downstream and upstream of the window.

**Analysis of psoralen-crosslinked chimeras**

We downloaded SPLASH sequencing reads from Poly-A enriched libraries of lymphoblastoid cell lines (LCL), human embryonic stem cells (ES), and retinoic acid (RA) differentiated cells^11^ from the Sequence Read Archive (accession number PRJNA318958). To map the reads, we first created a reference FASTA file containing nucleotide sequences spanning full gene lengths (ENSEMBL 105) and the 10 rRNA/spacer sequences. Then, we mapped the sequencing reads to the reference sequences using the bwa mem aligner^12^ with the following options: -a -c 1000000 (exhaustively report all alignments). According to the original pipeline, we collapsed reads that mapped to identical locations with identical alignment scores to aggressively remove PCR duplicates^11^. Finally, we extracted and annotated chimeric rRNA/spacer-mRNA and mRNA-mRNA reads requiring exact and correctly oriented matching of the pyknon in each portion of the chimeric read.
