## Supplementary material for "All ribosomal RNAs and 45S spacers from humans to worms are packed with organism-specific motifs whose other copies are found predominantly in numerous nervous system genes including many associated with human disorders": Supp. Figure S1

>rRNA-5S|1|-|228634869|228634989|.GRCh38

GTCTACGgccataccaccctgaaCGCGCCCGATCTCGTCtgatctcggaagctaagcagggtcGGgcctggttagtacttggatgggagacCGCCTGGGAATACCGGGTGCTGTAGGCTTT

>45S_rRNA.ETS1.GRCh38

[illegible]

>45S_rRNA.18S|21|+|8436876|8438744|.GRCh38

[illegible]

>45S_rRNA.ITS1.GRCh38

[illegible]

>45S_rRNA.5p8S|21|+|8439822|8439978|.GRCh38

CGACTCTTAGCGGTGGATCACTCGGCTCGTGCGTCGATGAAGAACGCAGCTAGCTGCGAGAATTAATGTGAATTGCAGGACACATTGATCATCGACACTTCGAACGCacttgcggccccgggttcctcccggggCTACGCCTGTCTGAGCGTCGCTT

>45S_rRNA.ITS2.GRCh38

[illegible]

>45S_rRNA.28S|21|+|8441146|8446211|.GRCh38

[illegible]

>45S_rRNA.ETS2.GRCh38

[illegible]

>rRNA-12S|MT|+|648|1601|.GRCh38

[illegible]

>rRNA-16S|MT|+|1671|3229|.GRCh38

[illegible]
