## Supplementary figures and images for "All ribosomal RNAs and 45S spacers from humans to worms are packed with organism-specific motifs whose other copies are found predominantly in numerous nervous system genes including many associated with human disorders"

### Supp. Figure S3

#### Supp. Figure S3

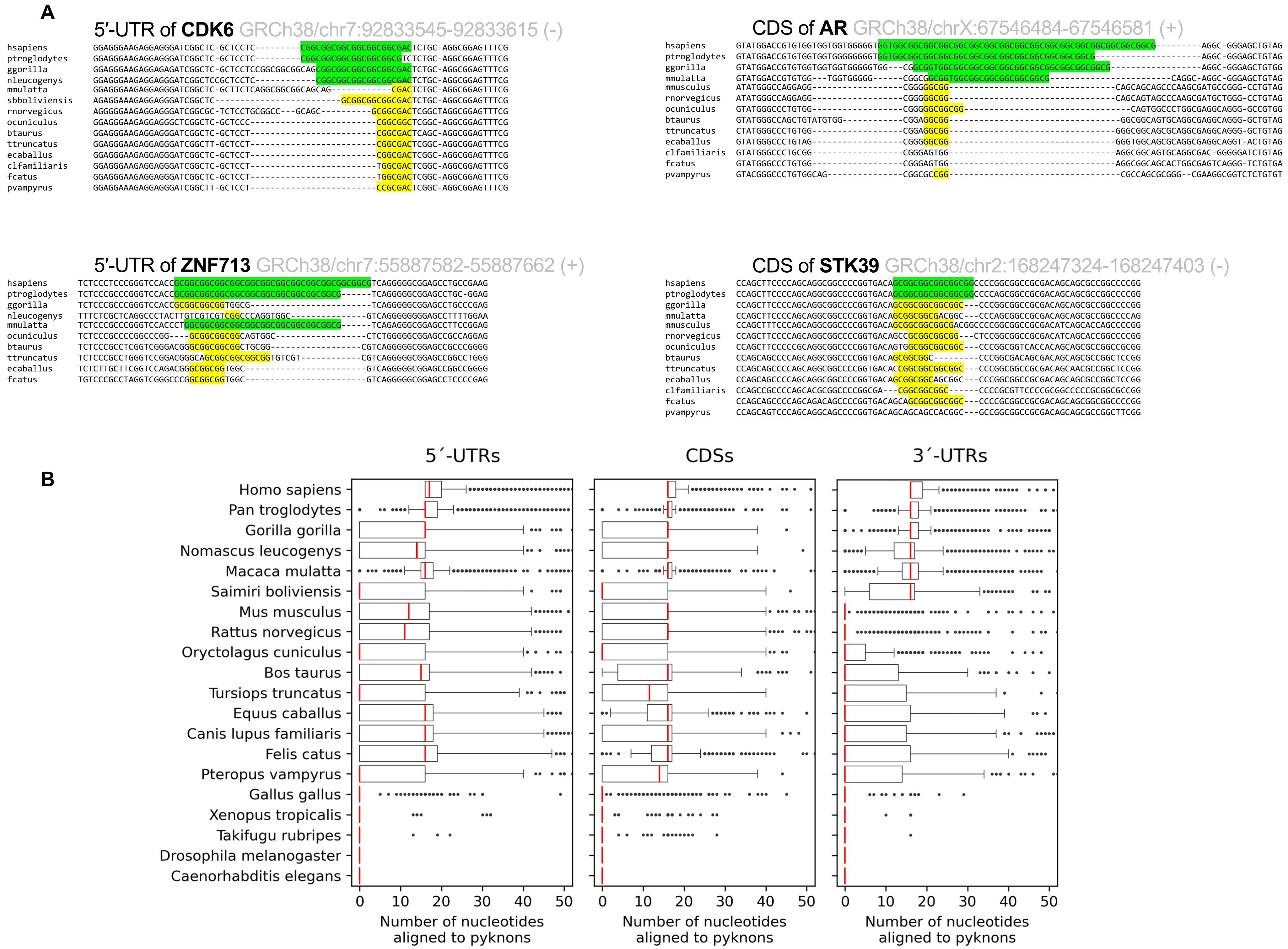

### Supp. Figure S4

#### Supp. Figure S4

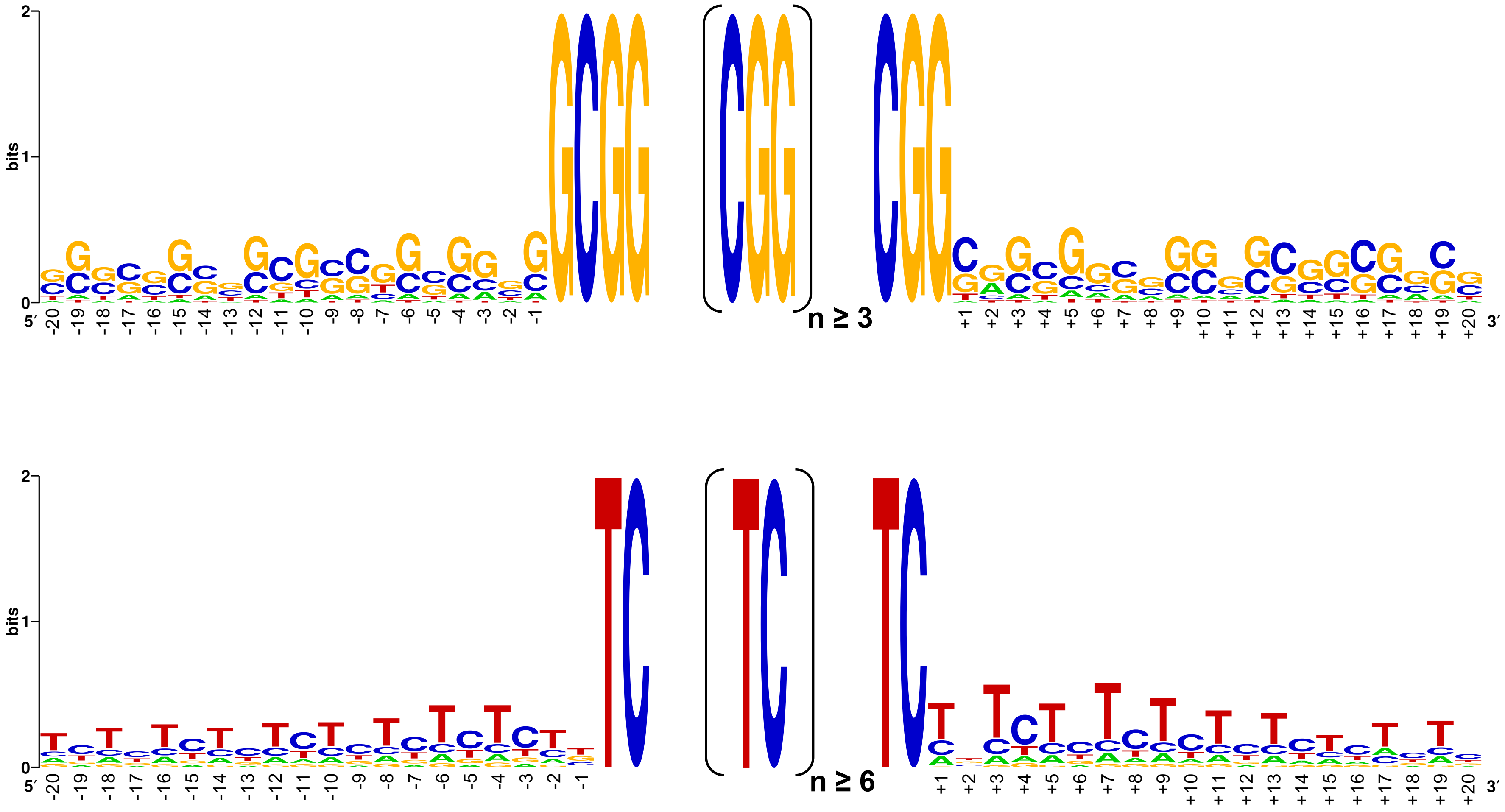

### Supp. Figure S5

Supp. Figure S5

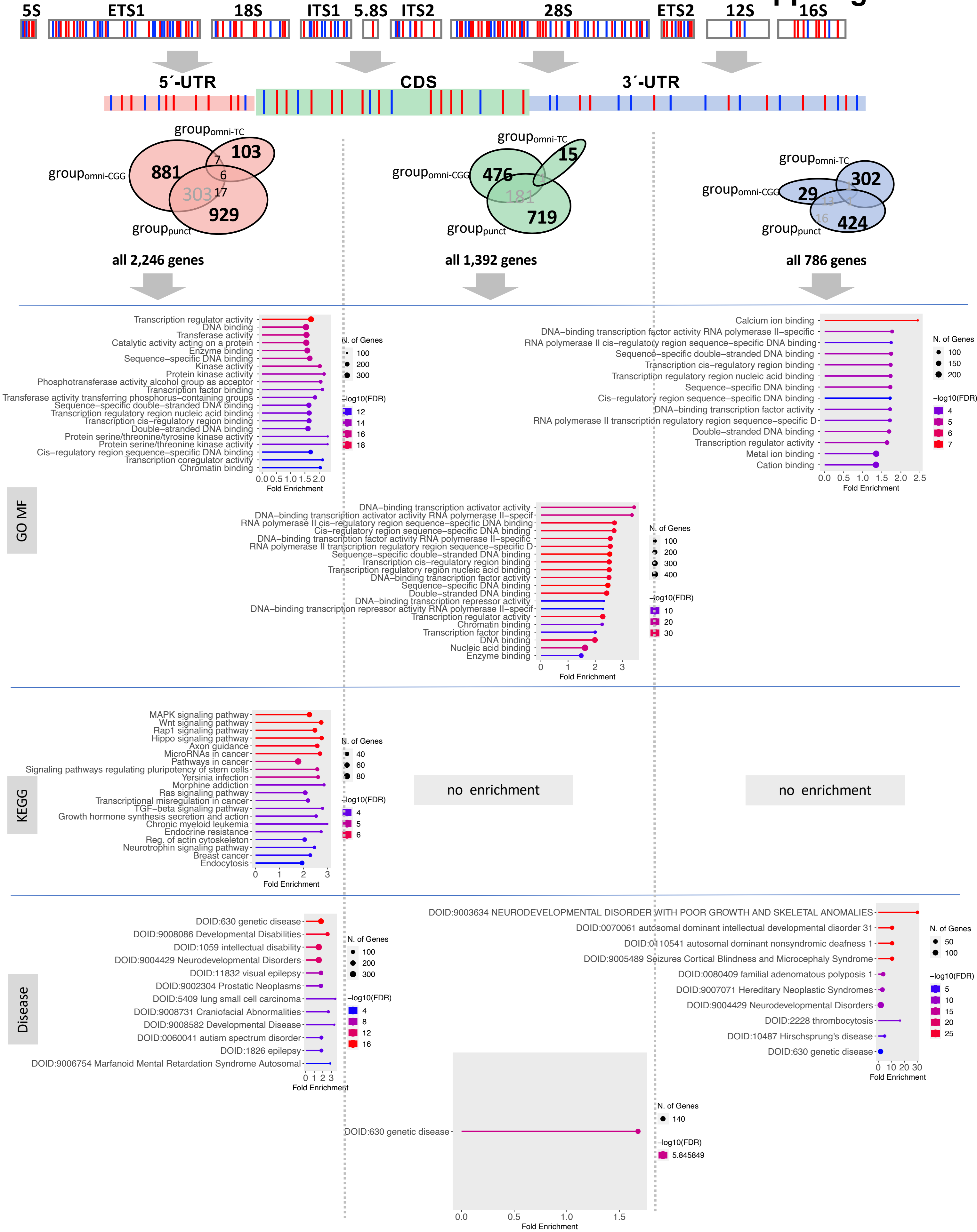

### Supp. Figure S6

#### Supp. Figure S6

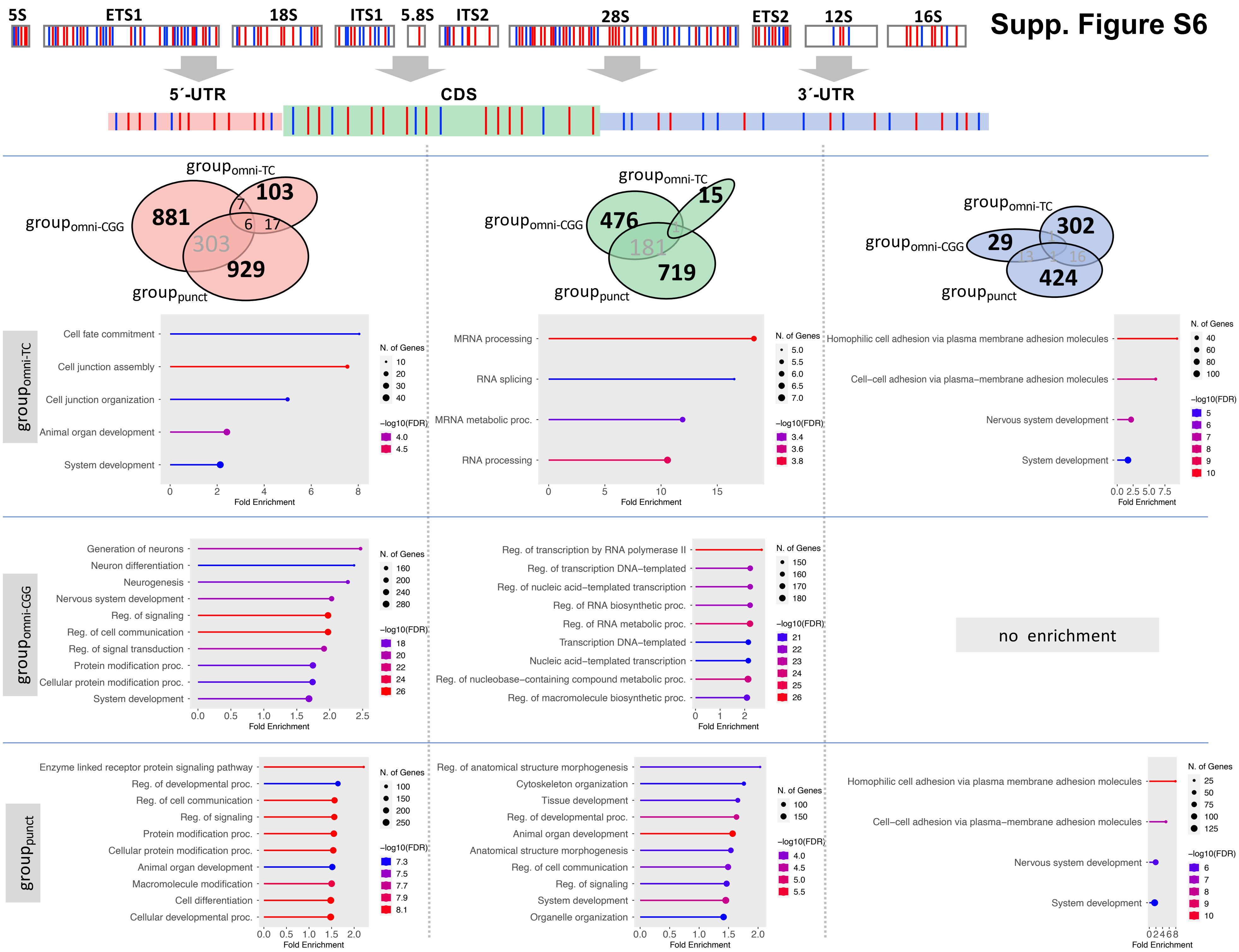

### Supp. Figure S7

#### Supp. Figure S7

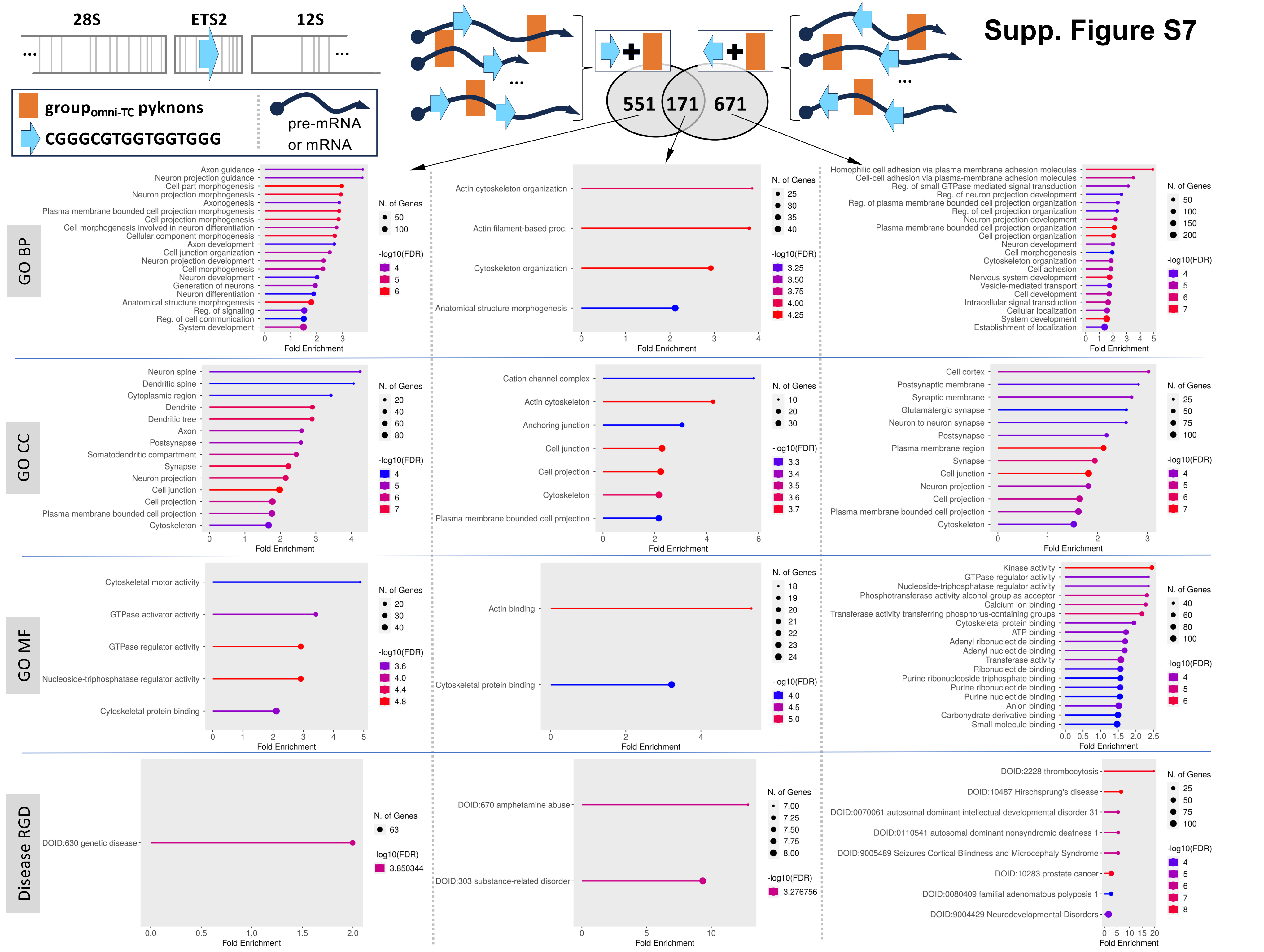

### Supp. Figure S8

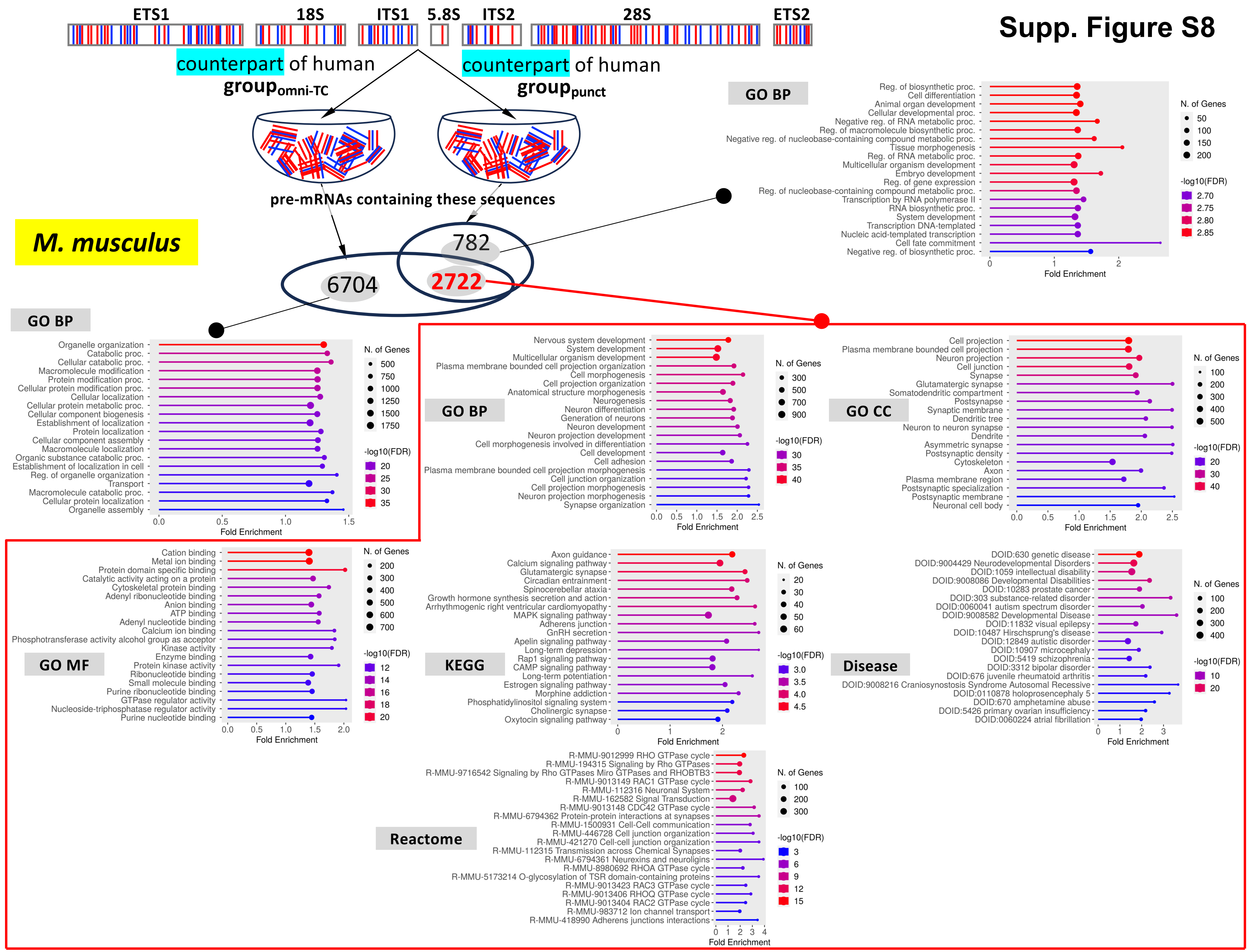

### Supp. Figure S9

#### Supp. Figure S9

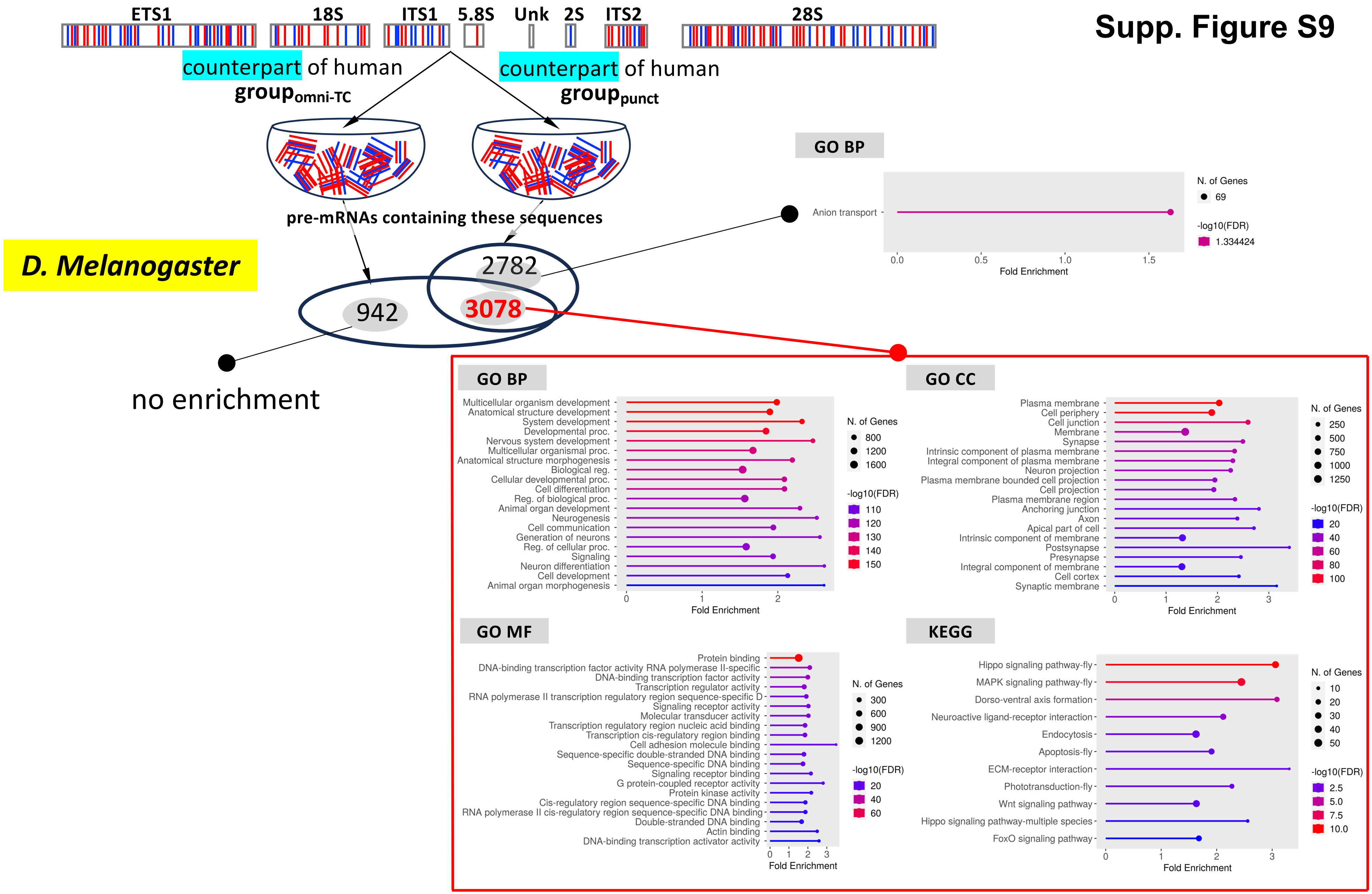

### Supp. Figure S10

#### Supp. Figure S10

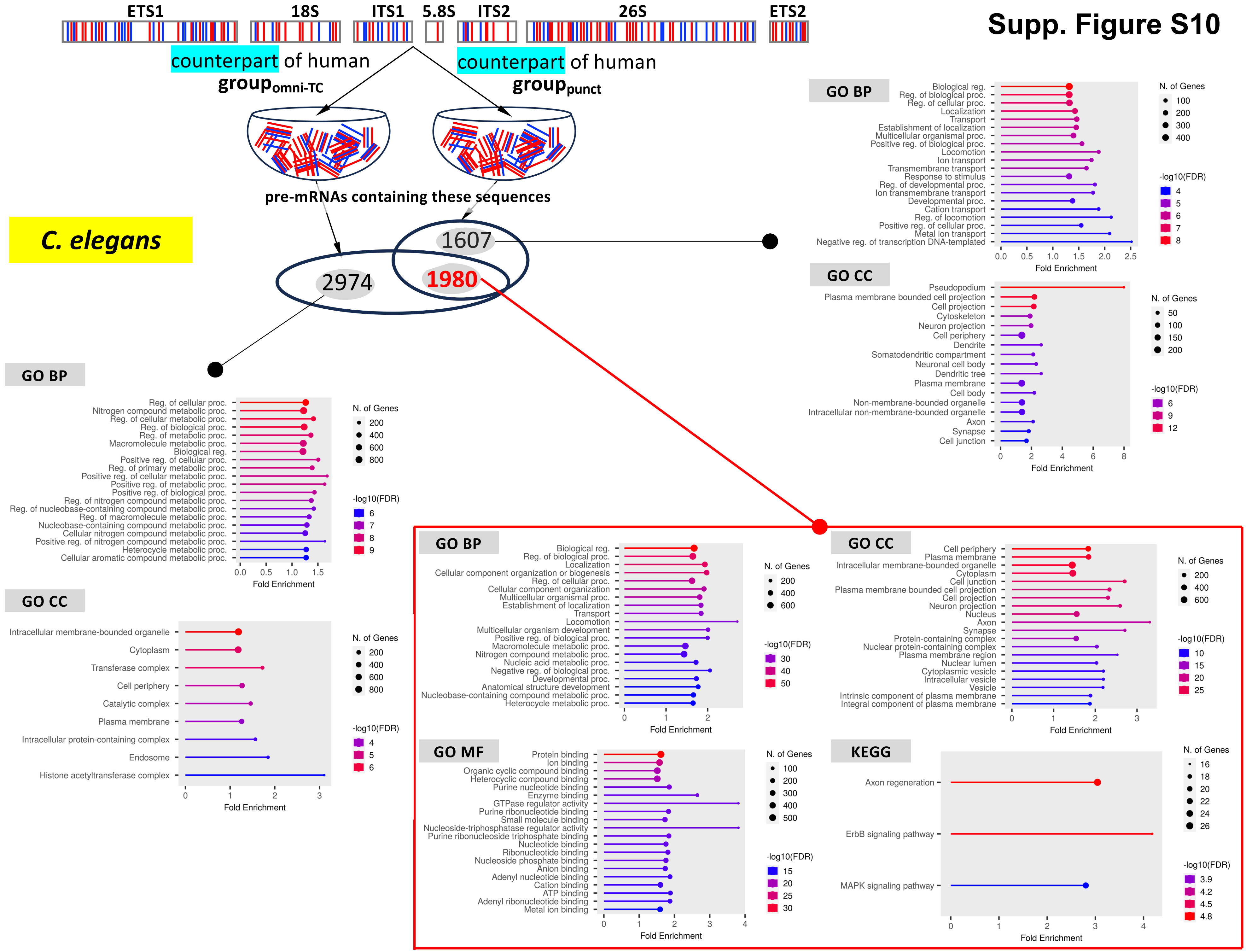
